# Retinotopic coding organizes the interaction between internally and externally oriented brain networks

**DOI:** 10.1101/2024.09.25.615084

**Authors:** Adam Steel, Peter A. Angeli, Caroline E. Robertson

## Abstract

The human brain seamlessly integrates internally generated thoughts with incoming sensory information, yet the large-scale networks that support these functions -- the (Default Network, DN) and external (Dorsal Attention Network, dATN) -- are traditionally viewed as functionally independent. This raises a crucial question: how does the brain integrate information across these seemingly non-interactive systems? Here, using densely sampled 7T fMRI, individualized resting-state parcellations, and voxel-wise population-receptive-field mapping, we show that these internal/external networks are more interlocked than previously thought. Spontaneous DN and dATN activity during rest is uncorrelated at the network level. However, voxel-scale functional coupling across networks is shaped by the latent visual field preferences of individual voxels in each network, as measured during independent retinotopic mapping. Voxels that share visual field preferences exhibit stronger spontaneous coupling than those with divergent preferences. These retinotopically-specific interactions are bivalent: DN voxels with negative (suppressive) visual response amplitudes are anticorrelated with matched (positive) dATN voxels, while those DN voxels with positive response amplitudes are positively correlated. Thus, distinct subpopulations of visually-tuned DN voxels participate in spatially-specific interactions with the dATN. Further, retinotopic coding is intrinsic to the DN, persisting even during periods when the DN signal is elevated. These findings reveal a latent, voxel-level architecture of retinotopically-grounded interactions between the DN and dATN. Taken together, our results suggest that retinotopic coding underpins the dynamic coordination of perception and thought in the human brain.

## Main Text

A fundamental goal of neuroscience is to understand how the coordinated activity of distributed brain networks gives rise to cognition (1–9). Central to this aim is understanding principles that govern interactions among networks with distinct computational roles. In particular, it is unclear how networks involved in externally-oriented attention (e.g., processing sensory input) interact with those that support internally-oriented attention (e.g., introspection and memory) (10–14). Understanding this interplay is crucial, because the dynamic exchange between internal and external representations underlies how we perceive, remember, and navigate our world -- yet the ’common language’ that enables communication between these systems remains unclear.

Historically, the Default Network (DN) and Dorsal Attention Network (dATN)—prototypical networks for internal and external attention, respectively—have been viewed as functionally competitive (10–12, 14, 15). Classic neuroimaging studies show that externally oriented tasks activate the dATN and deactivate the DN (10, 13, 14, 16–18), while internally oriented tasks produce the opposite pattern (6, 19–23). This dissociation is also observed at rest, where DN and dATN activity is typically independent or anti-correlated (11, 12, 24, 25), depending on the application of global signal regression(11, 26). Such findings have led to the prevailing view that DN/dATN antagonism supports selective attention to either internal or external information, minimizing interference between perception and memory (10, 12, 27, 28). Yet, this framework leaves open a fundamental question: if these networks are functionally dissociated, how does the brain integrate perceptual and mnemonic information when both are required – for example during anticipatory saccades or memory-guided attention?

Recent studies of the voxel-level coding properties within these large-scale networks offer insight into this question. Although the DN is traditionally associated with abstract or semantic coding(2, 28–30), emerging evidence suggests that it also exhibits a neural code typically associated with externally-oriented visual processing: coding of retinotopic stimulus position(31–34). Notably, retinotopic coding in the DN differs from that observed in traditional visually responsive areas, including the dATN. While visual stimulation elicits position-dependent *increases* in neural activity in traditional visual areas, many DN voxels often show position-dependent *decreases* in activity(31, 33, 34). This voxel-level pattern suggests that the DN’s global deactivation during externally-oriented tasks may not reflect a simple disengagement from external input, but rather a structured, spatially-tuned response to visual stimulation.

What purpose might the DN’s voxel-level retinotopic code serve? We and others have proposed that the visual code in the DN might play a functional role in structuring mnemonic-perceptual interactions (31, 32, 34, 35). Recent work provides critical support for this theory, revealing that scene-selective memory regions near the DN(36) exhibit voxel-level retinotopic opponency with scene-selective visual areas (34). Specifically, the inverted retinotopic code in mnemonic regions is functionally tethered to the positive code in perceptual regions: during perception, visual stimuli at specific field locations activate perceptual voxels while suppressing memory voxels tuned to the same location, a pattern that reverses during memory retrieval (34). These results indicate that visual field location can coordinate dynamic interactions between specialized perceptual and mnemonic areas at the cortical apex, raising the intriguing hypothesis that retinotopic grounding might support perceptual-mnemonic interactions more broadly across the brain. Yet, whether retinotopic coding serves as a more general organizing principle for interactions across large-scale networks remains unknown.

Here, we tested the hypothesis that global interactions across large-scale internally- and externally-oriented brain networks are structured at the voxel level by a shared retinotopic code^35^. This framework challenges the traditional view of internally- and externally-oriented brain networks as globally antagonistic, proposing instead that voxel-level retinotopic tuning enables functional coupling and shared information processing between them. To evaluate this hypothesis, we leveraged a high-resolution 7T fMRI dataset(37), individualized network identification(3, 4, 8), and voxel-wise modeling(38) to test three core predictions. First, voxel-wise retinotopic coding should be robustly present within individuals’ DN and dATN networks, but with divergent tuning polarities: positive retinotopic responses should dominate the dATN, while DN should contain a significant proportion of retinotopic voxels with negative response amplitudes to visual stimulation. Second, we predicted that spontaneous DN–dATN interactions during resting-state fMRI would be structured by the retinotopic properties of individual voxels. Specifically, functional coupling should be stronger between voxels that share visual field preferences across networks, revealing that retinotopic position-specific responses form a latent architecture to scaffold cross-network interactions even in the absence of overt task demands. Third, we predicted that these retinotopically grounded interactions would support both coupling driven by the DN and coupling driven by dATN, suggesting that shared spatial tuning enables dynamic, bidirectional interactions. Together, these findings would position retinotopy as a unifying framework for brain-wide information processing and for the coordination of internal and external attentional dynamics.

### Retinotopic coding in internally and externally oriented networks

To investigate the role of retinotopic coding in structuring interactions between internally- and externally-oriented brain networks, we analyzed 7T fMRI data from the Natural Scenes Dataset(37), acquired from the same densely-sampled participants during both resting-state and retinotopic mapping sessions (Fig. 1A-B).

**Figure 1.**
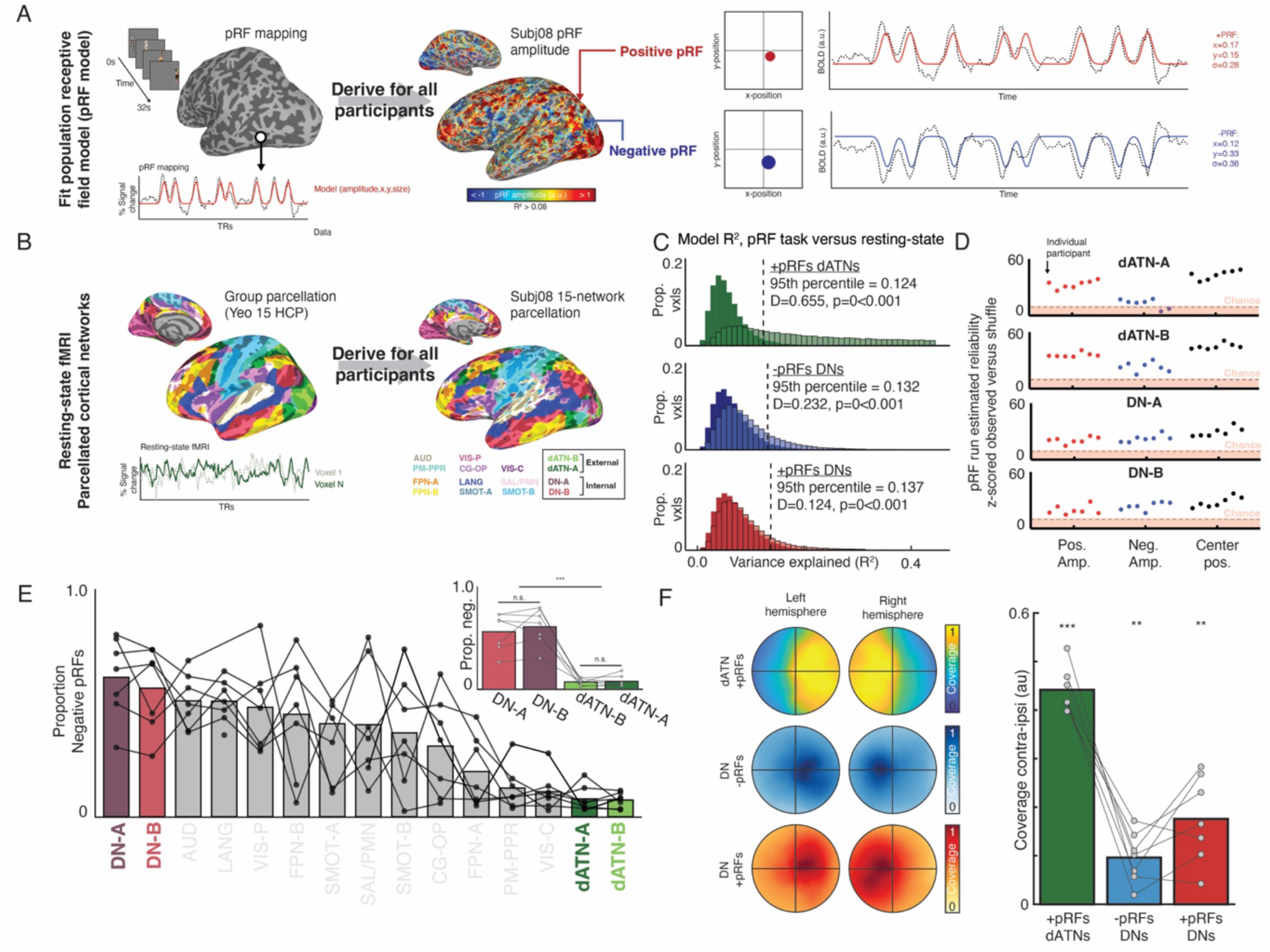
Inversion of retinotopic coding between externally- and internally- oriented brain networks. A. Population receptive field (pRF) modeling with fMRI. We established visual field preferences for each voxel using pRF modelling(38). Voxels with positive BOLD responses to the visual stimulus are classified as positive pRFs (+pRFs), and those with negative BOLD responses are classified as negative pRFs (-pRFs). B. Individualized resting-state network parcellation. Resting-state fMRI was collected in all participants (N=7; 34-102 minutes per participant) and used to derive individualized cortical network parcellations. Parcellations were generated using the multi-session hierarchical Bayesian modelling approach with the Yeo 15 HCP atlas as a prior(8, 42). C. PRF model variance explained is greater when fit to the pRF mapping task versus resting-state fMRI data for negative and positive pRFs in the DN and dATN. Histograms show voxel-wise variance explained for all pRFs fit to rest (dark) and the pRF modeling task fMRI data (light). All distributions were significantly shifted rightward for data fit to the pRF modeling task (dATN pRFs: K-S test: D^-^=0.655, p<0.001; Negative DN pRFs: D^-^=0.232, p<0.001; Positive DN pRFs: D^-^=0.124, p<0.001). Dashed lines indicate the 95^th^ percentile of R^2^ within the null distribution used; we chose the threshold for visually-defined voxels for subsequent analyses (R^2^ > 0.14) to exceed these values. D. DN and dATN pRF amplitude and center location estimates are reliable. To assess reliability of pRF amplitude estimates, we compared voxel-wise signed amplitude by computing the dice coefficients of binarized pRF amplitude maps (i.e., positive or negative pRFs) between pairs of pRF modelling runs. To estimate center position, we calculated the distance between pairs of pRF centers iteratively between pRF modeling runs. For both metrics, chance was estimated by performing the same computation on a random sampling of voxels drawn from both pRFs and non-pRFs (5000 iterations for each metric). In all subjects, for dATN positive and DN positive and negative pRFs, pRF amplitude and center position exceeded what would be expected by chance (z-scored Dice/distance values > 2.9). Although only dATN positive pRFs were included in subsequent analyses, the dATN pRF amplitude was not reliable above chance in two participants; this suggests that negative amplitude pRFs are not stable in all cortical networks. E. The internally-oriented DN contains a larger proportion of negative pRFs than the externally-oriented dATN. Proportion of negative pRFs (of total pRFs that survive thresholding in that network) in all cortical networks are shown. Inset. DN-A and DN-B contain more negative pRFs than the dATN (t(6)=7.94, p<0.001), and there is no difference in the proportion of negative pRFs among sub-networks. F. DN and dATN pRFs networks show contralateral bias. Coverage plots across participants for positive dATN pRFs and positive and negative DN pRFs. Bar plots show the difference in coverage between contra- versus ipsi-lateral visual fields for each pRF type. Bar plots show mean across participants, connected points represent individual participant data.

We first identified individualized large-scale networks for each participant using their resting-state fMRI data (8, 39) (Fig. 1B, S1). Resting-state data were preprocessed using independent components analysis (ICA) with manual noise component selection(40, 41), and no global signal regression was performed(26). Networks were identified using a multi-session hierarchical Bayesian modeling approach(8, 39, 42), which leverages repeated data runs to estimate within participant variation and derive stable parcellations for 15 cortical networks, including the Default Networks A-B and the Dorsal Attention Networks A-B (Fig. 1B). Unless otherwise noted, we combined the Default Networks A-B and the Dorsal Attention Networks A-B to constitute our internal and externally oriented networks, respectively (hereafter, DN and dATN).

Next, we determined the visual field preferences of all voxels within these individualized networks using a voxel-wise population receptive field (pRF) modeling approach(38, 43) to fit each participant’s BOLD activity during retinotopic mapping(38) (Fig 1A, S2). We fit the data using a gaussian model (implemented in AFNI) that allow for both positive and negative signed amplitudes.

We considered any voxel with >14% variance explained by our pRF model to be exhibiting a retinotopic code, and hereafter we refer to these voxels as “pRFs”. We determined this threshold by defining the noise floor of our retinotopic model in the DN and dATN by comparing task-derived model fits to a control null distribution derived from resting-state fMRI data (Fig. 1C; distribution of voxel-wise variance explained by task versus null, K-S tests: D^-^s>0.124, ps<0.001). Two pRF features were of primary interest: 1) center position, which estimates the preferred location in visual space that drives a given voxel’s response (x,y), and 2) amplitude of the voxel’s response (positive or negative) (Fig. 1A). Both positive and negative pRF amplitude was reliable in the DN for all subjects (all z-scored Dice coefficients > 2.9, Fig. 1D). In the dATN, positive amplitude pRFs were reliable in all participants, and negative amplitude pRFs (which constituted a small proportion of the overall pRFs in this network) were reliable in 5/7 participants. For the remainder of the paper, we only consider positive pRFs in the dATN. Importantly, pRF center position was highly reproducible across runs of pRF data in the dATN and DN in all subjects (Fig. 1D).

Next, we compared the retinotopic properties of voxels within the DN and dATN(11, 12, 18). Across participants, we observed a larger proportion of retinotopic voxels in the dATN as compared to the DN (43% of dATN voxels, 9% of DN voxels) consistent with their role as externally- and internally-oriented networks, respectively. Further, the amplitude of retinotopic responses (i.e. whether a stimulus in their preferred visual field location evoked a positive or a negative BOLD response) differed significantly between these two networks (Fig 1E). Consistent with prior reports, negative voxel-wise response amplitudes were clustered within the DN and in similar locations across participants (31, 34), evidenced by a consensus map of pRF sign (Fig. S3). On average, more than half (59%) of all pRFs in the DNs were inverted (i.e., had a negative BOLD response to visual stimulation in their population receptive field, -pRFs). In contrast, just 8% of the pRFs in the dATN were -pRFs (i.e., the vast majority of the voxels had positive BOLD responses to visual stimulation in their receptive field, +pRFs). The difference in negative pRF proportion between the dATNs and DNs was significant (t(6)=7.94, p<0.001; Fig. 1E, inset), and there was no difference among the sub-networks of each system (dATNs: t(6)=0.24, p=0.82; DNs: t(6)=0.89, p=0.41). This distinction is particularly remarkable given the proximity of dATN and DN clusters in posterior cerebral cortex. Like visual areas, both DN and dATN pRFs exhibited a contralateral visual field bias, as we would predict from prior literature (31, 33, 34) (positive dATN pRFs: t(6)=26.0, p<0.001; negative DN pRFs: t(6)=4.86, p=0.003; positive DN pRFs: t(6)=5.20, p=0.002) (Fig. 1F; see Fig. S4 for additional retinotopic properties). Together, these results show that individualized DNs have voxel-wise responses that exhibit visual coding -- the necessary foundation for assessing whether the voxel-wise retinotopic code structures global interactions between the internally- and externally-oriented networks.

### Visual responsiveness stratifies DN-dATN resting-state interactions

Next, we examined whether the retinotopic response properties of individual voxels (estimated during the retinotopic mapping sessions) predict their functional coupling across DN-dATN during task-absent, spontaneous activity (during the resting state scans). Specifically, we tested whether voxel-level visual response classes—non-retinotopic voxels, positively tuned pRFs (+pRFs), and negatively tuned pRFs (–pRFs)—predict the strength and direction of coupling between the DN and dATN. Such a finding would reveal a stable, voxel-level organizational principle underlying the global dissociation that is hypothesized to exist between the DN and dATN(11–13, 15, 24, 25).

Across resting state runs, global activity of the DN and dATN was independent (i.e., not correlated) after accounting for variance across all cortical networks using partial correlation(44) (mean correlation = 0. 0.031±0.13 s.d., t(6) = 0.6287, p=0.55; Fig. 2A). Crucially, however, this global relationship masked a striking stratification based on DN voxel responsiveness (Fig. 2B). Non-retinotopic DN voxels (i.e., pRF model R^2^<0.14) were not significantly correlated with the dATN (mean correlation = 0.014±0.135, t(6) = 0.26, p = 0.79). In contrast, DN voxels that responded positively to visual stimulation (DN positive pRFs, +pRFs) had a positive correlation with the dATN (mean correlation = 0.284±0.152, t(6) = 4.96, p = 0.0025), while DN voxels with systematic negative responses to visual stimulation (DN negative pRFs, -pRFs) were anti-correlated with the dATN (mean correlation = -0.21±0.149, t(6) = -3.75, p = 0.0094). This relationship was further strengthened by adopting more conservative R^2^ thresholds up to 0.30 to define the pRF population, despite the overall number of included voxels decreasing, suggesting that this effect is not driven by false-positive voxels at the edge of our threshold criteria (Fig. S5).

**Figure 2.**
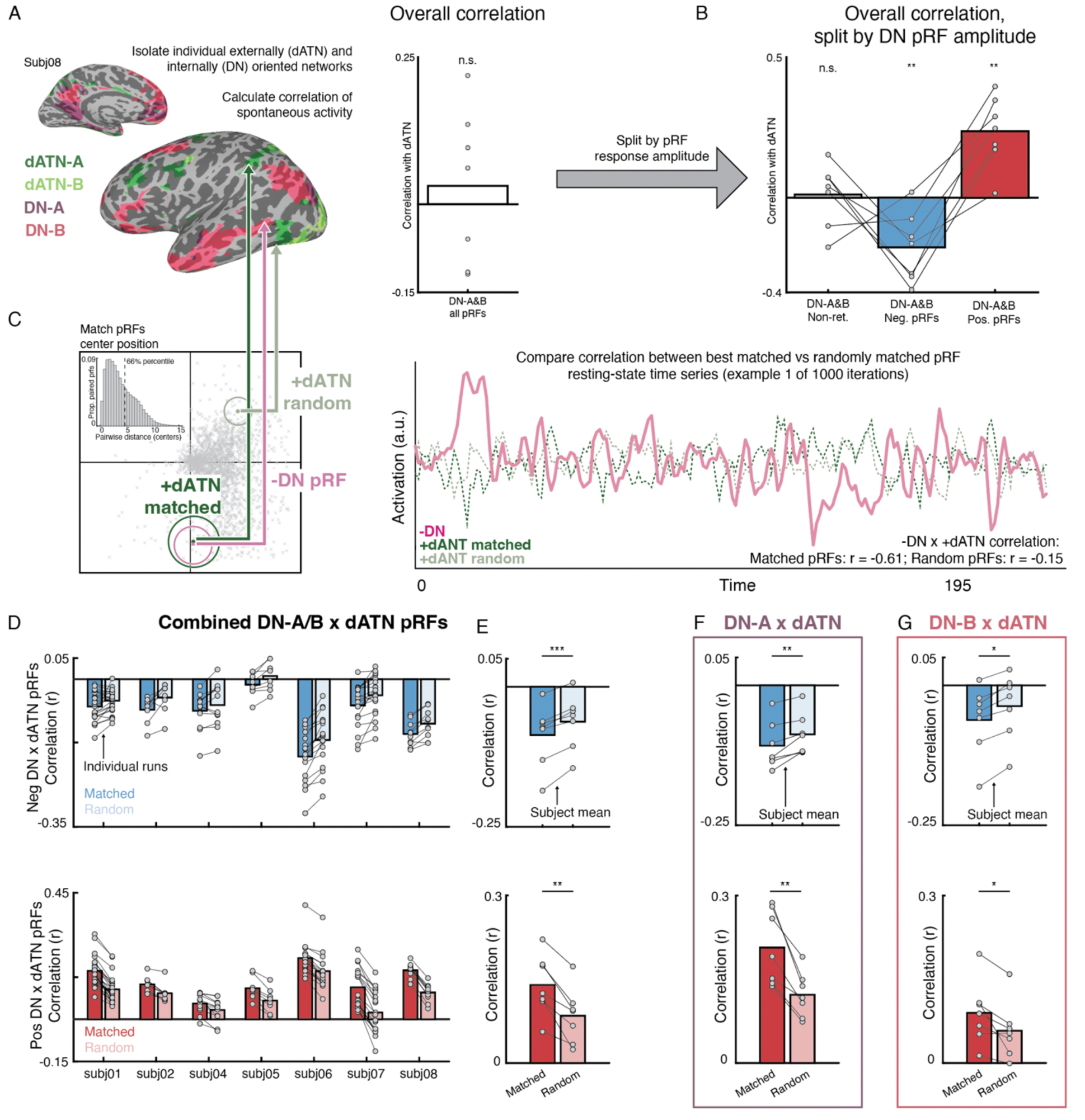
Retinotopic coding organizes spontaneous interaction between internally and externally oriented brain networks. A. Overall activity of the dorsal attention networks (dATN-A/B) and default networks (DN-A/B) were independent at rest. One individual’s dATN and DN are shown. Time series from all voxels in the DN and dATN subnetworks were averaged together, and we calculated the partial correlation(44) of these networks while accounting for variance of the other 11 networks within the group-prior parcellation(1, 8). The dATN and DN network correlation was not significantly different from zero, suggesting the activity of these networks is largely independent. Bar plot shows the average partial correlation across participants, each data point is the average of all runs for a single participant. B. Interaction between the DN and dATN differs by visual field preference of DN voxels. Splitting the DN voxels by visual responsiveness revealed distinct patterns of resting-state correlation. DN +pRFs had a positive correlation with the dATN (mean correlation = 0.284±0.152, t(6) = 4.96, p = 0.0025), while non-retinotopic DN voxels (i.e., pRF model R2 < 0.14) were not significantly correlated with the dATN (mean correlation = 0.014±0.135, t(6) = 0.26, p = 0.79). On the other hand, DN -pRFs were anti-correlated with the dATN (mean correlation = - 0.21±0.149, t(6) = -3.75, p = 0.0094). C. Determining spatially-matched pRFs in dATNs and DNs. We assessed the influence of retinotopic coding on the interaction between internally- and externally-oriented brain areas’ spontaneous activity during resting-state fMRI, by comparing the correlation in activation between pRFs in these networks that represent similar (vs. different) regions of visual space. For each DN -pRF, we established the top 10 closest positive dATN pRFs’ centers based on their x and y position (“matched”), and correlated the DN pRF time series with the average timeseries of these matched dATN pRFs. We compared these correlation values with the correlation of the DN pRF timeseries paired with 10 randomly selected pRFs from the 33% furthest (1000 iterations per pRF). We computed this for all DN pRFs for each all runs in each subject. Inset shows the pairwise distances between all pRFs in all participants, dotted line shows the 66%ile. D. Negative (upper) and positive (lower) DN pRFs show retinotopic specific interaction with the dATN. Each set of bars depicts the average correlation of the matched and randomly paired pRFs between the DN and dATN for each participant, with each run shown as connected points. All participants showed greater correlation (either positive or negative) for matched compared with randomly paired pRFs. E. Data shown averaging runs within each participant. For both positive and negative pRFs, the correlation between matched compared with randomly matched pRFs was significantly greater (consistent with their amplitude, ps<0.01). F, G. show the correlation between positive and negative pRFs in the DN with the dATN, split across the sub-networks of the DN-A and B. Both DN-A and B show a significant impact of matching, indicating that both subnetworks interact with the dATN via a retinotopic code (ps<0.05). *: p<0.05, **: p<0.01, ***:p<0.001

This result shows that voxel-level visual response profiles shape DN-dATN coupling during spontaneous resting-state activity. Specifically, the DN and dATN activation is independent during rest. However, at the voxel-level, specific sub-groups of DN voxels have distinct coupling patterns with the dATN that depends on the valence of voxels’ visual responses. DN and dATN voxels with positive visual responses show a positive relationship during rest, and a notable subset of DN voxels with negative visual responses display the canonical opponency with dATN voxels. This suggests that retinotopic coding may be a mechanism that enables visual information to be exchanged between these large-scale brain systems.

### Shared visual field preferences structure functional coupling among visually-responsive DN and dATN voxels

We next asked whether functional coupling between DN and dATN voxels is modulated by the visual field preferences of retinotopic voxels — that is, whether their pRFs are centered on similar versus different visual field locations (i.e., their retinotopic alignment). We hypothesized that DN–dATN interactions would be stronger for voxel pairs with shared visual field preferences. To test this, we quantified the retinotopic alignment of voxel-wise pRFs across the DN and dATN.

Because we only considered positive pRFs in the dATN, hereafter we refer to these as “dATN pRFs.”

For each pRF in the DN (+pRFs and –pRFs), we computed the Euclidean distance between its receptive-field center (x, y) and those of all dATN pRFs (Fig. 2C). We then identified the 10 closest dATN pRFs (“Matched” set) and averaged their resting-state time series. As a comparison group, we repeated this procedure using 10 randomly selected, poorly matched dATN pRFs (“Random” set; see Methods). This procedure was repeated 1000 times for each DN voxel across resting-state runs.

Matched dATN pRFs were distributed throughout the network, spanning posterior retinotopic cortex to anterior regions, including prefrontal cortex (Fig. S6). Crucially, the spatial dispersion (i.e., the distance between the pRFs across the cortex) of matched and randomly-matched pRFs did not significantly differ (matched versus random – DN -pRFs x dATN pRFs: t(6)=0.38, p=0.71; DN +pRFs x dATN pRFs: t(6)=0.67, p=0.52), which suggests that spatial proximity is not a confounding factor.

If DN-dATN interactions are scaffolded by a retinotopic code, spatially matched pRFs should exhibit stronger functional coupling than randomly-matched pRFs. Consistent with this prediction, after partialling out variance from DN +pRFs to isolate the specific co-fluctuation between DN - pRFs with dATN pRFs (34, 44), DN -pRFs remained anticorrelated with dATN pRFs in nearly every resting-state run (Fig. 2D). Critically, matched pRFs were significantly more anticorrelated compared with randomly-matched pRFs (linear mixed-effects model with random intercepts for subject and run: F(1,194) = 105.7, p < 0.001; coefficient = –0.023 ± 0.002 s.d.). This effect was evident when run-level correlations were averaged within each participant (7/7 participants r_matched_ < r_random_; t(6)=6.9, p<0.001, Fig.2E) and could not be explained by signal quality differences, as matched and random pRF sets did not differ in tSNR (t(6) = 1.01, p = 0.353). Finally, to rule out any possible influences from vascular stealing (i.e. the shunting of blood into active tissue from nearby regions), we repeated the matching analysis after excluding any voxels within a 3mm radius of a major vessel (Fig. S7; see Methods). Both matching effects remained after excluding vascularly susceptible voxels (+DN x dATN: t(6) = 6.054; p < 0.001; -DN x dATN: t(6) = -5.0448; p < 0.01). This is strong evidence that the retinotopic code underpins the opponent interaction between negative DN and positive dATN pRFs.

Voxel-wise retinotopic alignment scaffolded the interaction between positive DN and dATN pRFs as well. After controlling for DN -pRFs, DN +pRFs were positively correlated with dATN pRFs (Fig. 2D). Retinotopic alignment again strengthened the interaction: matched correlations exceeded random correlations in a mixed-effects model (F(1,194)= 406.5, p < 0.001; coefficient = +0.050 ± 0.003 s.d.) and for every individual participant (7 / 7; t(6) = 5.7, p = 0.001) (Fig. 2E).

Together, these results demonstrate that shared retinotopic alignment enhances functional coupling between DN and dATN voxels. Voxels tuned to the same region of visual space show stronger spontaneous co-fluctuation—whether positively or negatively coupled—than those with non-overlapping spatial preferences. This suggests that a shared spatial code helps organize spontaneous interactions between large-scale internal and external attention networks during rest.

### Subnetwork Specificity within the DN

Recent work has suggested that the subnetworks of the DN, DN-A and DN-B, support different cognitive processes: episodic projection and theory of mind, respectively (8, 45–47). Indeed, DN-A is particularly sensitive to the process of scene construction(47), closely linked to visual imagery, suggesting that dATN pRFs would show greater interaction with visually-responsive voxels in DN-A compared with DN-B. Consistent with this hypothesis, spontaneous correlations with dATN pRFs were significantly stronger for DN-A than DN-B voxels (collapsed across positive and negative pRFs: t(6) = 4.80, p = 0.003). Crucially, however, voxel-scale retinotopic alignment structured interactions between the DN and the dATN for both DN sub-networks (Fig. 2F-G; DN-A: -pRFs t(6) = –4.56, p = 0.004; +pRFs t(6) = 5.65, p = 0.0013; DN-B: -pRFs t(6) = –7.49, p < 0.001; +pRFs t(6) = 2.92, p = 0.027). The strength of this effect did not significantly differ between DN-A and DN-B (t(6) = 1.51, p = 0.182). Thus, although DN-A is more strongly globally coupled with dATN, retinotopic preferences of individual voxels structure functional interactions across both DN subnetworks equally.

In addition, using the same matched-vs-random pRF procedure, we found that resting activity between the lateral place-memory area (LPMA) and the adjacent occipital place area (OPA) was significantly more correlated for retinotopically matched than random voxel pairs (t(6)=3.01, p=0.024; SI). This shows that retinotopic scaffolding within domain-specific systems extends beyond perceptual and memory tasks (34), organizing spontaneous activity within these systems as well. These findings build on and extend our previous work (34) and others(48).

### Retinotopic coding is intrinsic to internally-oriented areas

Our findings thus far highlight a role for retinotopic coding in structuring spontaneous interactions between a subset of voxels across large-scale internally- and externally-oriented brain networks. However, a key mechanistic question remains: is retinotopic coding an intrinsic property of the DN, or does it only emerge in response to input from sensory systems and lower-level networks like the dATN? One possibility is that DN-driven modulation of dATN activity reflects a global, non-selective suppression, with retinotopic coding in the DN arising only through input from the dATN. Alternatively, retinotopic coding may be intrinsic to the DN itself – such that signals projecting from the DN to the dATN are retinotopically-specific, allowing the DN to shape fine-grained spatial representations within the dATN. Disambiguating these possibilities is essential for determining whether the retinotopic scaffold constitutes a general mechanism for cross-network communication, independent of task demands or cognitive domain (e.g., memory versus perception).

To address this, we again turned to participants’ resting-state fMRI data. Because participants are not engaged in any task, resting-state provides a neutral testbed for assessing spontaneous DN–dATN interactions. Moreover, experience sampling during rest has shown that attention naturally alternates between internal and external focus during rest (49, 50), and thus is likely to result in activation of both the DN and dATN, making it possible to assess whether retinotopic coding emerges primarily during periods of elevated DN or dATN activity.

We developed an event-triggered analytical approach to dissociate periods when activity of the retinotopic voxels within the dATN and DN was elevated. We refer to these periods when dATN or DN voxels’ activity was elevated as dATN-driven from DN-driven events, respectively. Specifically, we identified timepoints where spontaneous activity of individual DN or dATN pRFs exceeded the 99th percentile of their resting-state distribution. These high-activation time points were treated as neural “events” (Fig. 3A-C). To test whether these events produced retinotopically-specific responses in the target network, we compared peri-event activation in the target network’s most 10 closely matched pRFs (i.e., those with similar visual field preferences) to that in the 10 worst-matched (anti-matched) pRFs (Fig. 3A). We used the worst matched pRFs for this analysis because the large number of pRFs made the random matching procedure impractical. All analyses were conducted separately for positive and negative pRF populations within the DN. This design enabled us to test two competing predictions: if retinotopy in the DN only reflects input from more canonically perceptual networks like the dATN, we should only see retinotopically specific coupling during dATN-driven events. If, however, the DN possesses an intrinsic retinotopic scaffold, we should observe retinotopic specificity during both dATN- and DN-driven events. A windowed correlation analysis produced similar results (Fig. S8).

**Figure 3.**
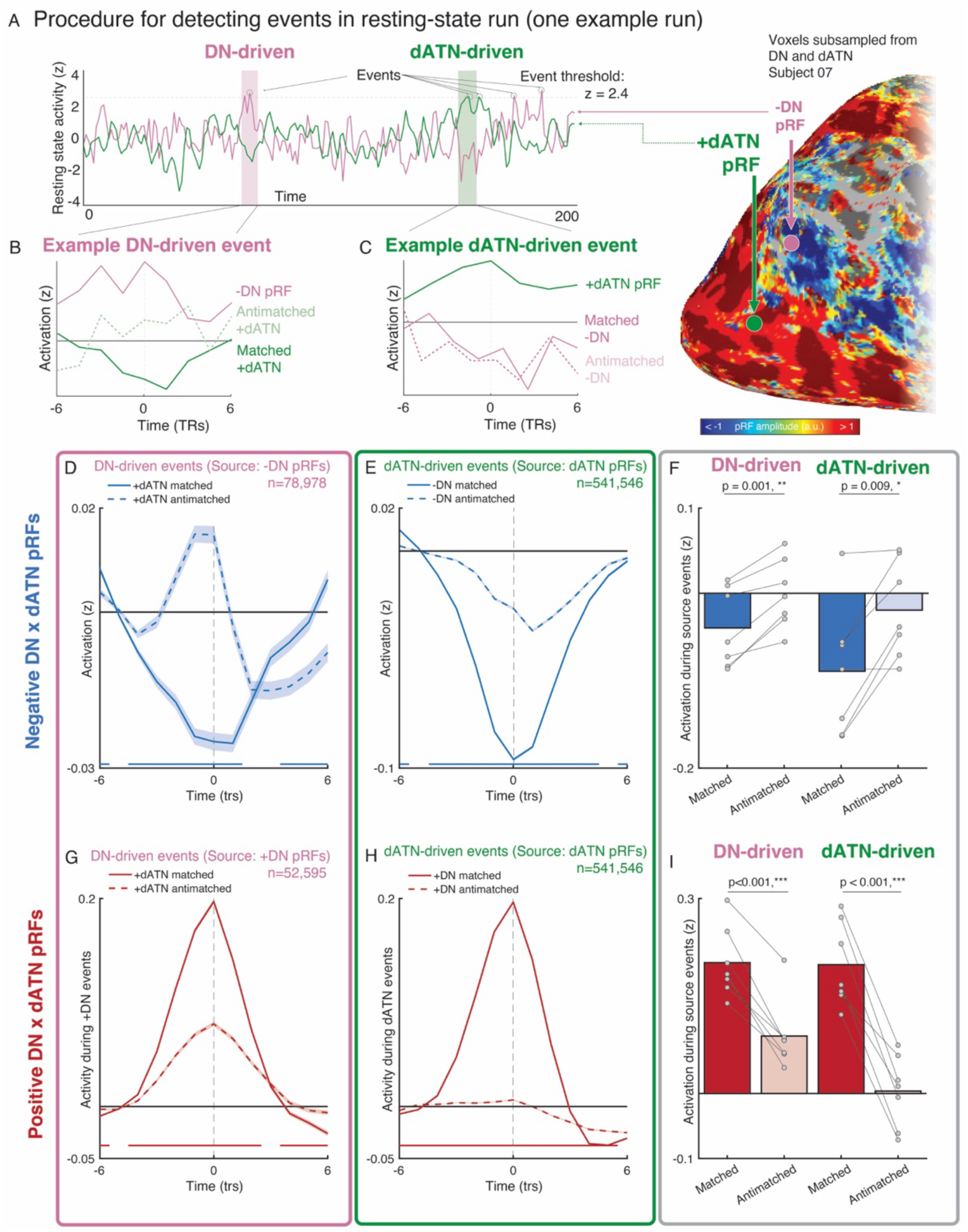
DN-driven vs. dATN-driven neural events detected in spontaneous resting-state dynamics show evidence for retinotopically-specific suppression. A. Event detection and analysis procedure and example events from a single resting-state run. To detect events, we extracted the time series from each pRF in the source regions (positive or negative pRFs in the DN; positive pRFs in the dATN) and isolated time points where the z-scored time series exceeded 2.4 s.d. (99th percentile). We then examined the activity of 10 best matched and 10 worst matched (anti-matched) pRFs from the target region in this peri-event time frame (6 TRs (8 s) before and after the event). B. Example DN-driven event in the representative time series shows elevated source pRF activity (DN -pRF) and corresponding target area activity (dATN +pRF). C. Same as B, but for dATN-driven activity with the event detected in a dATN +pRF. D, E. Evidence for retinotopically-specific suppression of ongoing activity during DN-driven events detected in DN -pRFs (D) and dATN-driven events detected in dATN +pRFs (E). D. Time series show the peri-event activity of matched and anti-matched dATN +pRFs averaged across all participants for events detected in DN -pRFs (N=78,978 events). E. Time series shows the peri-event activity in DN -pRFs during events detected in dATN +pRFs (N=541,546). F. For DN -pRFs paired with dATN +pRFs, target area shows retinotopically-specific suppression of activity for both DN-driven and dATN-driven events. Bars show the average activation at event time of the target areas’ matched and anti-matched pRFs for each participant. Activity was lower in matched compared with anti-matched pRFs for DN-driven and dATN-driven events (ts>3.8, ps<0.009). G, H. Evidence for retinotopically-specific excitation of ongoing activity during events DN-driven events detected in DN +pRFs (G) and dATN-driven events detected in dATN +pRFs (H). G. Grand average time series for matched and anti-matched dATN +pRFs during events detected in DN +pRFs for all participants. H. Grand average time series for matched and anti-matched DN +pRFs during events detected in dATN +pRFs. I. For DN +pRFs paired with dATN +pRFs, target area shows retinotopically-specific excitation for both DN-driven and dATN-driven events. Bars show the average activation at event time of the target areas’ matched and anti-matched pRFs for each participant. Activity was higher in matched compared with anti-matched pRFs for DN-driven and dATN-driven events (ts>8.75, ps<0.001). Interestingly, the activity of anti-matched pRFs was greater during DN-driven compared with dATN-driven events (t(6)=4.92, p=0.0026), suggesting that input from DN +pRFs to the dATN +pRFs may be more global and less retinotopically-targeted compared with dATN-driven signaling between these networks. For time series plots, solid bars along the x-axis indicate periods with a significant difference between matched and anti-matched pRF activity within the target area. *: p<0.05, **: p<0.01, ***:p<0.001. -DN: Negative DN pRFs; +DN: positive DN pRFs; +dATN: Positive dATN pRFs

Across all participants, we detected 78,978 events in DN -pRFs, 52,595 events in DN +pRFs, and 541,546 events in dATN pRFs. dATN-driven events were more frequent than either DN-driven type (dATN vs. negative DN: t(6)=15.19, p<0.001; dATN vs. positive DN: t(6)=25.63, p<0.001), but there was no difference between DN-driven event types (t(6)=2.03, p=0.18). On average, participants exhibited 2082±535 (mean±SD) negative DN, 1799±400 positive DN, and 3102±602 dATN events per run. Each pRF showed 0–4 events per run (Fig. S9). More than 90% of timepoints involved <10 simultaneous events, suggesting that individual pRF-level correlation could be isolated from global fluctuations in regional activity, making this approach suitable for evaluating distinct interactions at the individual voxel level.

We found a robust effect of retinotopic alignment (i.e., matched versus anti-matched) on pRF activation during both dATN- and DN-driven interactions (Fig. 3D-I). For DN -pRFs, a linear mixed-effects model (factors: DN-driven/dATN-driven and retinotopic alignment; subject as random effect) revealed a main effect of match status (F(1,24)=15.3, p=0.0006) and no interaction between match status and direction (F(1,24)=0.0748, p=0.40). During DN-driven events in DN - pRFs, matched dATN pRFs showed lower activity than anti-matched (t(6)=5.46, p=0.001) and were significantly deactivated relative to the pre-event baseline. The same was true of DN -pRFs during dATN-driven events: we observed lower activation for matched vs. anti-matched pRFs (t(6)=3.80, p=0.009). These results are consistent with a retinotopically-specific interaction between dATN +pRFs and DN -pRFs during both DN-driven and dATN-driven events detected in spontaneous activity during resting-state fMRI.

The interaction between DN +pRFs and dATN pRFs also showed evidence of retinotopic coding during both dATN- and DN-driven events, but these effects were modulated by a direction × match interaction (F(1,24)=12.84, p=0.002). Both DN-driven (t(6) = 8.76, p < 0.001) and dATN-driven (t(6) = 9.14, p < 0.001) events evoked strong activation in the target network (dATN or DN, respectively). However, anti-matched pRFs were more active during DN-driven compared to dATN-driven events (t(6) = 4.92, p = 0.0026), suggesting that signals from positive DN pRFs to dATN may be less retinotopically specific than incoming signals from the dATN. These results align with prior findings showing decreased spatial precision in visual representations during memory recall compared to perception(51, 52).

Taken together, these results support the view that retinotopic coding serves as a general scaffold for DN-dATN interactions. This alignment shapes spontaneous coupling during both periods of DN-driven and dATN-driven activity, even in the absence of external tasks. Because the effect persists without sensory input, it likely reflects an intrinsic feature of the DN activity rather than a transient result of drive from sensory input.

## Discussion

Despite extensive evidence that internally- and externally-oriented brain networks – including the default network (DN) and dorsal attention network (dATN) – are critical for human cognition(1–8, 11–13), the coding principles that enable these distributed systems to interact have remained unclear(30, 53–57). Here, we show that DN–dATN coupling is retinotopically grounded at the voxel scale: the visual field biases of individual voxels scaffold bidirectional interactions between these networks, even during spontaneous dynamics in the absence of visual task demands. Analyses of transient elevated activity in either the DN or dATN confirm that this retinotopic opponency shapes both dATN-driven signals to the DN and DN-driven signals to the dATN, indicating that the retinotopic code is intrinsic to both internal and external networks. These findings offer a multi-scale account of neural communication, in which interactions among sub-populations of voxels with shared tuning preferences are nested within macro-scale network dynamics. Nesting multiple neural codes might enable ongoing computations within a larger brain system (e.g., attending to internal mental states within the DN during memory recall), while simultaneously allowing for the sharing of fine-grained representations across brain systems (34). Here we show this latent, voxel-scale interaction persists even without structured behavioral demands, opening the opportunity to study these fine-scale interactions across brain networks in resting-state data (58).

Resolving the coding principles of the DN is critical given its central roles in spatial and episodic memory, social cognition, and executive function(13, 59–64). Classic views of systems-level brain organization posit regions at the apex of the cortical hierarchy, like the DN, likely have abstract, amodal neural codes, in contrast to sensory codes–like retinotopy–that manifest in earlier areas of the cortical hierarchy (2, 29, 30). Our findings and others challenge this view. Instead of shedding the retinotopic code at the cortical apex, the DN retains a retinotopic code(31, 33) and functionally deploys this code to structure interactions with the externally-oriented dATN. Thus, a code once thought to be exclusive to early visual cortex and visually-oriented tasks plays a widespread role in shaping brain activity across the cortical hierarchy—even at rest.

Establishing this voxel-level structure within the DN and dATN reframes a long-standing debate that has cast these networks as globally competitive or functionally independent(11, 23, 27, 28).

These views have stemmed from macro-scale observations of reciprocal activation patterns during external versus internal task demands(6, 10, 13, 14, 16–23), and resting-state analyses that reveal zero or negative correlations between their mean time series(11, 12, 24, 25). However, by focusing on global, network-level activity across voxels, these approaches have obscured a latent, voxel-level structure to DN-dATN functional coupling. Specifically, we find that DN–dATN coupling is not independent. Instead, two subpopulations of voxels within the DN contribute to a retinotopically-grounded cross-network interaction that is stratified by their voxel-level visual selectivity. One population of DN voxels is positively tuned to visual stimulation, and positively covaries with dATN voxels that share their visual field preference. In contrast, DN voxels that are negatively tuned to visual stimulation negatively covary with retinotopically matched dATN voxels as a function visual field preference. When these voxel-level tuning preferences are not taken into account, global cross-network opponency or independence(11, 12) reinforces the misconception that the networks support mutually exclusive cognitive modes. By revealing a shared retinotopic scaffold, our findings reconcile why DN and dATN can appear independent in coarse analyses yet remain functionally integrated, offering a new framework for understanding how internal and external attention are coordinated in the human brain.

Importantly, there are key differences in the nature of retinotopic coding between the DN and visual areas, including the dATN. First, retinotopy in the DN is predominantly negative in valence—DN voxels more often show spatially-specific suppression in response to visual stimulation(31, 33, 34), unlike the positive spatially-specifical response that is typically observed in visual areas. Second, the DN lacks the map-like retinotopic organization seen in early visual cortex. In visual areas, retinotopic coding is not only the foundational code, but also the dominant topographic organizing principle across the cortical surface, where voxels that are proximal on the cortical surface tend to share visual selectivity for nearby regions of visual space(65). Prior work has emphasized the *visual response bias* in regions where voxel-wise retinotopic responses lack a map-like organization(35); overall, the DN does exhibit this kind of bias. However, our results show that the voxel-scale activity underpinning this bias reflects the latent connectivity of those voxels. Thus, we adopt the term “retinotopic coding”, because this voxel-scale coding scheme exists without a map-like organization on the cortical surface. For example, rodent and bat head direction cells are not laid out in a literal ring, but the population code of these neurons forms a ring manifold(66, 67). Similarly, in humans, retrosplenial cortex shows head-direction coding during navigation tasks despite lacking topographic organization of head-direction responses(68). Analogously, the DN may implement a retinotopic code that is distributed across the network in a format that does not spatially mirror the retina. That is, the DN may use a fine-scale retinotopic code without a retinotopic map – the representational structure can be preserved even in the absence of spatially ordered cortical layouts.

What function might the bivalent (positive and negative) voxel-scale retinotopic alignment across these large-scale networks serve? Our previous work has identified a role for retinotopic-alignment in structuring the functional coupling of scene-selective mnemonic and perceptual brain areas(34, 48) located near the dATN and DN, respectively (36, 69). Near to the DN, scene-selective memory areas exhibit a negative retinotopic code, while functionally-paired scene-selective perceptual regions exhibit a positive retinotopic code(34). This bivalent retinotopic code structures the mutual activity of these mnemonic and perceptual areas: scene-recall activates scene-memory voxels, it but suppresses voxels in scene-perception areas that are tuned to matched visual locations, a pattern that reverses during scene perception(34). The current work significantly extends this observation beyond perceptual and mnemonic task-states -- identifying widespread retinotopic alignment across large-scale networks with presumed different orientations (internal vs. external) even during spontaneous activity at rest, in the absence of tasks.

As to the functional role of the bivalent retinotopic alignment across internal and externally-oriented large-scale networks, we propose two complementary possibilities. One possibility is that the bivalent (positive and negative) subpopulations of DN pRFs may serve as a neural mechanism for maintaining separate channels for perceptual and mnemonic information(54), preventing mnemonic signals from being interpreted as or interfering with sensory signals. A second possibility is that bivalent coding across the DN and dATN could be a mechanism for signaling mnemonic predictions via top-down inhibitory processes(34, 70). Specifically, when the DN generates predictions about incoming sensory information based on memory, the inverted retinotopic code could act as a suppressive signal that sharpens tuning in perceptual areas and enhances signal-to-noise for expected stimuli(71, 72). These mechanisms may work in concert: the inverted code simultaneously maintains separate channels for memory/perception within the DN, while enabling precise top-down predictive signaling across DN-dATN. Future work that combines high-resolution recordings with behavior will be essential for testing how these mechanisms jointly support constructive operations such as memory-based prediction during visually guided navigation.

Relatedly, the DN is considered a transmodal hub for cortical processing, where disparate sensory and motor processes converge (59, 73) The DN’s position at the cortical apex implies connections with and influence over unimodal cortical areas. However, the mechanism linking unimodal and transmodal networks had been unknown. Prior work posited that sensory coding in transmodal areas might serve this function (31, 35), and our data provide direct empirical support for this account: specific visually-responsive voxels provide an input/output interface bridging perceptual and memory systems. This complements work delineating specific affective and effective subregions within the DN that link the DN to other brain areas (74). Thus, while the DN may be “distant from input”(28), these data suggest that it is not disengaged from sensory processing.

Retinotopic alignment between the DN and dATN was clear in both subnetworks of the DN, DN-A and DN-B(3, 8, 20). This is surprising given their distinct functional profiles: DN-A has been more directly linked to visually grounded memory processes, such as episodic projection and scene construction(8, 20, 47), and spatially overlaps with regions previously shown to exhibit retinotopy and engage in scene memory and perception(69). In contrast, DN-B is more often associated with processes that are not obviously tied to vision: theory of mind and affective tasks, such as interpreting false beliefs or representing physical and emotional pain (8, 20). Why might the DN-B show retinotopically-grounded coupling with perceptual networks? Retinotopic coding might still be a useful format for transforming abstract social representations into sensory-grounded signals to inform perceptual predictions (e.g., anticipating changes in facial expression) or behavioral decisions (e.g., making eye movements). Future work might consider direct manipulation of social perceptual processes to assess the impact of social tasks on retinotopic coding between DN-B and dATN.

Moments of high spontaneous activity originating in the DN co-occurred with suppression of retinotopically-matched dATN pRFs, suggesting that retinotopy functions as a native coding format in the DN for targeted modulation of perceptual cortex. This result aligns with other recent studies suggesting an important role of the DN in shaping visual processing. For example, ongoing pre-stimulus DN activity has been shown to influence sensitivity during near-threshold visual object recognition(75). Other work has shown that subareas of the DN encode kinematic of spatial exploration in rodents(76), and represent the visuospatial context that is associated in memory with a perceived scene(34, 36, 69, 77, 78). Relatedly, DN activity reflects semantic-level attentional priorities during visual search(79). Taken together, these findings show that, rather than being disengaged during visual tasks, the DN actively shapes responses in perceptually-oriented cortex in a spatially-precise manner. In this sense, retinotopic coding is part of the DN’s “native language”. Taken together, these findings prompt a reevaluation of the DN’s role in perceptual processing, and the extent to which roles in “internally- and externally-oriented attention” adequately captures the DN and dATN’s role in cognition.

Is the DN a “visual” network? One possible interpretation of retinotopic coding in the DN is that the DN directly and obligatorily represents retinal position in addition to more abstract features(38, 65). Under this hypothesis, the retinotopic code may indicate that the information represented in the DN is, in part, sensory. Alternatively, the apparent visual coding may simply represent an underlying connectivity structure(80), which enables associative and semantic information that the DN directly represents(30, 81, 82) to be effectively transferred to sensory areas of the brain. Under this alternative hypothesis, the information represented in the DN is not itself sensory in nature; the retinotopic code in the DN simply serves as a “highway” between the abstract representations in the DN and sensory representations in the dATN. Notably, in either case the retinotopic scaffolding represents a latent structure linking high-level and low-level areas through consistent, spatially-organized interactions(80). This provides more evidence that low-level sensory codes are distributed across the brain, joining recent work showing latent somatotopic codes in regions of the brain that are not typically associated with somatosensation, including visual cortex(83). The wide and overlapping distribution of sensory codes across the brain may allow for multiplexing of complex information to structure cross-network interactions(35, 83). Further studies investigating this framework with diverse tasks and stimuli could reshape our understanding of how the brain processes and integrates information across different levels of cognitive complexity.

Our study and other recent studies of voxel-wise visual responses in traditionally non-visual areas (31, 33, 34, 48, 84–86) raise important questions about model selection and thresholding criteria. In canonical visual cortex, pRF analyses often rely on variance-explained thresholds to identify reliable voxel-wise model fits. However, it is less clear what R² criterion should be applied in non-visual regions, where responses may be weaker or more heterogeneous. To address this issue, we designed a method to determine a threshold for voxel-wise model fits (R^2^) empirically by leveraging resting-state fMRI data as a noise model. Crucially, pRF parameter estimates (center position and amplitude) for voxels that passed this empirical threshold were reliable across runs, suggesting this method robustly selects voxels with stable pRF properties. This empirical method may be used evaluate retinotopic responses across the brain in future studies (86).

Our study made use of the NSD, which included pRF modeling data to characterize visually-responsive cortex in a small group of individual participants. One limitation of this dataset is the small number of pRF task runs, which might lead to noisier model estimates. Our robust thresholding approach mitigates this concern. Furthermore, our primary results were based on voxel-wise pRF alignment within independently acquired resting-state fMRI data; noisy estimates of pRF parameters would obscure this relationship and make our results less likely. Relatedly, here we used a single gaussian model, consistent with prior work on negative visual responses in memory systems (31, 33, 34). However, other models of visual receptive fields might offer further insight into the DN’s visual responsiveness, such as double gaussian models of surround suppression (87) or compressive summation (88). Future studies might directly compare different visual models to further refine the computations underpinning visual responses in the DN.

In summary, our results show that a voxel-level retinotopic code organizes the spontaneous interactions between large-scale internally- and externally-oriented networks in the human brain – the DN and dATN. Even when global activity appears uncorrelated across DN-dATN, voxel-level visual-field tuning reveals a tightly interlocked, bidirectional functional dynamic that persists in the absence of external stimulation. This latent scaffold reconciles the apparent macroscale independence of internally- and externally- oriented systems with their evident functional need to cooperate, revealing that the DN is not divorced from perceptual networks, but modulates perceptual voxels through a common visuospatial reference frame. Taken together, by grounding large-scale network interactions in a shared, bivalent spatial code, retinotopy may offer a simple organizing principle for the brain’s coordination of thought and perception.

## Materials and Methods

### Dataset and Participants

The data analyzed here are part of the Natural Scenes Dataset (NSD), a large-scale precision 7T fMRI dataset consisting of high-resolution anatomical scans, retinotopic mapping, functional localizers for visual areas, and both task and resting-state fMRI data for eight adult participants. A full description of the dataset can be found in the original manuscript (37). Here, we detail the data, processing, and analysis steps relevant to the present work.

Participants (N = 8; 2 male, 6 female; ages 19–32) were scanned at the University of Minnesota over the course of 33-43 sessions. All participants had normal or correct to normal vision and no known neurological impairments. Informed consent was collected from all participants, and the study was approved by the University of Minnesota institutional review board. One subject (subj03) was excluded from resting-state analyses for having insufficient resting-state runs that passed our quality metrics (see below).

For the present study, we focused on two components of the NSD protocol: retinotopic mapping and resting-state fMRI. All participants completed three runs of a sweeping bar retinotopic mapping task (5 mins each). Retinotopic mapping was conducted during Session 1, along with the functional localizer scans. This task is referred to as “RETBAR” in NSD documentation.

All participants completed at least 14 runs of a resting-state fixation task (range: 14-34 runs; 5 mins each). Resting-state data runs were included in all NSD scan sessions after session 20 (i.e., session 21, 22, etc.). Each of these sessions included two 5-minute resting-state runs -- one at the start and one at the end of the session -- yielding 14 to 34 total resting-state runs per participant. During the second resting-state run, participants performed a deep breath in the first 5 seconds of scanning. To account for this, and to ensure the two resting-state runs were equated in length, we excluded the first 30 seconds of the resting-state data from all runs. One participant (subj03) was excluded from resting-state analyses due to having an insufficient number of usable runs after quality control. All remaining participants had at least 8 high-quality resting-state runs (mean: 14 ± 6.13 runs).

### MRI acquisition and processing

For this study we made use of the following data from the NSD: anatomical (T1 and FreeSurfer segmentation/reconstruction(89, 90)), functional regions of interest (ROIs), and minimally preprocessed retinotopy and resting-state time series. All analyses except network identification were conducted in original subject volume space, and data were projected on the surface for visualization purposes only. Network identification was performed on the fsaverage6 surface, then projected into subject specific volume space for quantification.

#### Anatomical data

Anatomical data was collected using a 3T Siemens Prisma scanner and a 32-channel head coil. We used the T1-weighted anatomical images provided at 0.8mm resolution as well as the registered output from FreeSurfer recon-all, aligned to the 1.8mm functional data. For venography, we used the provided time of flight (TOF) MR venography images with a resolution of 0.5mm^3^. For visualization purposes, we projected statistical and retinotopy data to the cortical surface using SUMA (91) from the AFNI software package(92).

#### Functional MRI data acquisition and processing

All analyses were conducted on 1.8mm isotropic resolution minimally processed runs of a sweeping-bar retinotopy task and resting-state fixation provided in the NSD.

##### Quality assessment

To ensure only high-quality resting-state data were included, we trimmed the first 25 TRs (40s) from each run (77), which left 4.25 min of resting-state data per run. We then used AFNI’s quality control assessment tool (APQC(93)) on the raw trimmed resting-state time series to assess the degree of motion in the resting-state scans. We excluded runs with greater than 1.8mm maximum displacement per run or 0.12mm framewise displacement from analysis, and assessed runs with greater than 1mm displacement or 0.10mm framewise displacement on a case-by-case basis(8). After exclusions, one participant (subj03) had only a single run that survived our criteria (4.25 minutes of data)), so we excluded them from our analyses. The remaining 7 participants had at least 8 resting-state runs (> 34 minutes of data; mean number of runs: 14±6.13 (sd), range: 8-24 runs).

##### ICA denoising

To further denoise the retinotopy and remaining resting-state data, we used manual ICA classification of signal and noise on the minimally-preprocessed time series(36, 40, 41). We used manual classification because automated tools perform poorly on the high spatial and temporal resolution data of the NSD(94). For each retinotopy and resting state run, we decomposed the data into independent spatial components and their associated temporal signals using ICA (FSL’s melodic(95, 96)). We then manually classified each component as signal or noise using the criteria established in(41). Noise signals were projected out of the data using fsl_regfilt(40).

We did not perform global signal regression(26). Data were normalized to percent signal change. A 2.5mm FWHM smooth was applied on the surface to resting-state fixation data used in functional connectivity network identification. The analysis of voxel-scale interactions performed on unsmoothed data.

#### Data analysis – Identifying Resting-State Networks

##### Individual-specific cerebral network estimation

In each participant, we identified a set of 15 distributed networks on the cerebral cortical surface using a multi-session hierarchical Bayesian model (MS-HBM) functional connectivity approach (8, 39). Briefly, the MS-HBM approach calculates a connectivity profile for every vertex on the cortical surface derived from the top 10% of correlations from that vertex to 1,175 ROIs uniformly disturbed across the cortical surface(1). We applied an expectation-maximization algorithm, initialized with a 15-network group-level spatial prior created by applying previous methods of network identification(1) on a portion of the HCP S900 data, to all the connectivity profiles from all participants. This algorithm then estimated each network’s group-level connectivity profile, along with parameters for subject-, session/run-, and region-level variability, which together were used to generate individual-specific, labeled parcellations.

The MS-HBM approach is well-suited to derive parcellations for the participants in the NSD, where resting-state data was collected across multiple days, because this method uses repeated resting-state runs for each participant to estimate within-participant variance. This leverages a strength of the NSD (the relatively large number of resting-state runs for each individual) and can account for differences in connectivity both within an individual (likely the result of confounding variables such as scanner variability, time of day, etc.), and between participants (reflecting potentially meaningful individual differences). Moreover, the MS-HBS incorporates a group spatial prior, further stabilizing the resulting network parcellations for each participant. This also ensures that all networks will be identified in all participants, while allowing for idiosyncratic topographic differences.

#### Data analysis – Retinotopic Mapping

##### Population receptive field (pRF) Modelling

The NSD retinotopy stimuli consist of a mosaic of faces, houses, and objects superimposed on pink noise that are revealed through a continuously drifting aperture. For our analysis, we considered only the bar stimulus time series, which is consistent with other studies investigating negative pRFs in high-level cortical areas (31, 33, 34). We did not consider the wedge/ring stimulus for any analyses.

After denoising the retinotopy data, we averaged the three retinotopy runs with the bar aperture together to form the final retinotopy time series. We performed population receptive field modeling using AFNI following the procedure described in (43). First, because the pRF stimulus in the NSD is continuous, we resampled the stimulus time series to the fMRI temporal resolution (TR = 1.333s). Next, we implemented AFNI’s pRF mapping procedure (3dNLfim). Given the position of the stimulus in the visual field at every time point, the model estimates the pRF parameters that yield the best fit to the data: pRF amplitude (positive, negative), pRF center location (x, y) and size (diameter of the pRF). Both Simplex and Powell optimization algorithms are used simultaneously to find the best time series/parameter sets (amplitude, x, y, size) by minimizing the least-squares error of the predicted time series with the acquired time series for each voxel. Relevant to the present work, the amplitude measure refers to the signed (positive or negative) degree of linear scaling applied to the pRF model, which reflects the sign of the neural response to visual stimulation of its receptive field.

##### Identifying Retinotopic Voxels – Empirically-derived R² thresholds

We developed a rigorous, data-driven approach to determine whether our pRF model fits truly reflect visual coding rather than physiological noise. To do this, we leveraged the resting-state fMRI data – where there is no dynamic visual stimulation -- to create a control (null) distribution for retinotopic model fits and establish a “noise floor” for each network (i.e., the DN and dATN). Then, we compared task-derived model fits to this rest-based null distribution to establish an empirical R² threshold for pRFs in each network.

Specifically, we fit the pRF model to the average timeseries across all resting-state runs for each participant. We then compared the resulting voxel-level R² distribution to that from the pRF mapping task data, pooled across participants. To test whether the model fits were significantly higher for the task vs. null distribution, we performed a one-tailed Kolmogorov-Smirnov test for each network on the respective cumulative distribution functions (H^0^: CDF^rest^(R^2^) ≤ CDF^task^(R^2^), H^a^: CDF^rest^(R^2^) > CDF^task^(R^2^)). These analyses were performed separately for the dATN and positive and negatively responsive voxels within the DN.

We defined a voxel as exhibiting a retinotopic code (hereafter referred to as a “pRF”) if its model fit exceeded 14% variance explained. This threshold was based on the 95th percentile of the null R² distributions, which yielded noise floors of R² = 0.124 (dATN), R² = 0.132 (DN-positive), and R² = 0.137 (DN-negative). We therefore set our global threshold for visually responsive voxels across all networks at R² > 0.14, as this value exceeded the noise floor calculated in each network separately.

##### Quantifying visual field coverage

Visual-field coverage (VFC) plots represent the sensitivity of an ROI across the visual field. We followed the procedure in Steel et al., 2024(34), which we have reproduced here. Individual participant VFC plots were first derived based pRF center, as well as pRF size and R^2^. These plots combine the best Gaussian receptive field model for each suprathreshold voxel within each ROI by calculating the max operator at each point in the visual field, defined as the maximum value from all pRFs within the ROI. The resulting coverage plot thus represents the maximum envelope of sensitivity across the visual field. Individual participant VFC plots were averaged across participants to create group-level coverage plots.

To compute the elevation biases, we calculated the mean pRF value (defined as the mean of the average max operator values in a specific portion of the visual-field coverage plot) in the contralateral upper visual field (UVF) and contralateral lower visual field (LVF) and computed the difference (UVF–LVF) for each participant, ROI and amplitude (+/−) separately. A positive value thus represents an upper visual-field bias, whereas a negative value represents a lower visual-field bias.

##### Reliability of pRF amplitude and center position estimates

We tested whether key pRF features of interest – amplitude (i.e., positive vs. negative) and center position (i.e., x,y) – were reliable across retinotopy runs. To assess the reliability of pRF amplitude, we iteratively compared the amplitude of significant pRFs from individual runs of pRF data. Specifically, for each retinotopy run, we fit the pRF model and then binarized any significant voxels (R² > 0.14) by amplitude. When testing positive-pRF reliability (in dATN and DN), positive voxels = 1; all others = 0. For negative-pRF reliability (in DN), significant negative voxels = 1. We then computed a three-run Dice coefficient: (dice 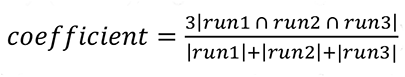). We compared this value against 5000 null Dice scores derived from the same number of voxels randomly sampled from all voxels, both significant and non-significant, in the ROI. For each participant, this resulted in 1 “observed” dice coefficient along with 5000 null values representing the distribution of dice coefficients expected by chance. The observed coefficient was z-scored against this null distribution for each participant.

We also evaluated center position (x,y) reliability. For every significant voxel, we calculated the Euclidean distance between its center estimates across the three run pairs (i.e. 1-2, 2-3, 1-3). As with amplitude, we generated a null distribution by repeating this computation on 5,000 random voxel sets (significant + non-significant). To establish significance of the true estimate, the observed estimate was z-scored against this null distribution for each participant.

#### Data Analysis – Retinotopically-Grounded Resting-state Analyses

##### Evaluating global functional coupling between cortical networks at rest

To assess global functional coupling between the default network (DN) and the dorsal attention network (dATN) at rest, we used a partial correlation analysis that accounted for variation associated with other cortical networks (44). For this and all subsequent analyses, each individual resting state run was z-scored in time. Prior to calculating the partial correlation between them, we first averaged the time series within each DN and dATN subnetwork.

##### Evaluating the Contributions of Voxel-Level Visual Response Valence to Cross-Network Coupling

To test whether DN–dATN coupling is modulated by the visual responsiveness (i.e., response valence) of DN voxels, we repeated this analysis separately using only: (i) sub-threshold/non-retinotopic voxels, (ii) positive pRFs, or (iii) negative pRFs from the DN -- while still partialing out the remaining networks. This allowed us to assess whether specific classes of visually responsive voxels in the DN differentially contribute to functional coupling with positive pRFs in the dATN.

##### Evaluating Voxel-Level Retinotopic Alignment in Cross-Network Coupling

To test whether functional coupling is shaped by the shared visual field preferences of DN-dATN voxels (i.e., retinotopic alignment), we compared resting-state correlations between pRF pairs across the DN and dATN that had matched (vs. unmatched) pRF centers.

To do this, for each pRF in the DN (positive or negative), we computed the Euclidean distance between its receptive-field center (x, y) and those of all dATN +pRFs. We focused on center position instead of pRF size because 1) size estimates are relatively unstable compared with center position(97), and 2) we lacked a clear prediction about how pRF size would be related between the two networks. For each DN voxel, we identified the 10 nearest dATN +pRFs based on center position—the “matched” set—and averaged their resting-state time series. As a null comparison, we defined a “random” set by sampling with replacement 10 dATN +pRFs from the most distant third of the match distribution (i.e., (the 1/3 furthest pRF pairs). We considered the bottom 33% because the overwhelming majority of pRFs constituted relatively equal “good” matches. This random sampling was repeated 1,000 times per DN voxel, generating a distribution of 1,000 random correlation values, which we then averaged to yield a single “random” correlation for each DN source pRF.

We calculated the correlation between each DN pRF’s resting-state time series and that of its 10 best-matched dATN pRFs (“matched” correlation). We averaged across the 10 best matches to (1) reflect the likely distributed input from multiple dATN sources and (2) improve signal-to-noise. To isolate pRF-specific effects, we partialed out the mean time series of the complementary DN pRF population from the signals of interest. For example, when analyzing correlations between – DN and +dATN, we treated +DN activity as a nuisance regressor (and vice versa)(34, 44). This provided a network-level analog to the whole-brain partial correlation described above (see Section: Correlation among cortical networks), allowing us to isolate the unique contribution of the pRFs of interest from overall activity in the region as well as to control for global fluctuations that cause widespread positive correlations like motion and attentional state(34, 44). All correlation values were Fisher-transformed, and we took the median value across voxels within each network to yield a single value per run. This procedure was repeated for all runs and all participants.

We then statistically compared Fisher-transformed correlations for matched vs. random pRF pairs using a linear mixed effects model, with match status (matched vs. random) as a fixed effect and subject and run as random effects. This model accounted for both between- and within-subject variation across session. For follow-up tests, we compared the average matched and random values across participants using a paired t-test.

##### Masking Vascularly Sensitive Voxels

To ensure that the above retinotopic alignment analyses were not the result of large-scale vascular features (e.g. vascular stealing), we created whole brain mask to remove any voxels within 3mm of a significant vessel. We identified and enhanced vessels within the 0.5mm TOF MR venography image using a Hessian-based multiscale vesselness filter (Matlab command *fibermetric*), then binarized the result based on a threshold calculated from a fit 2-Gaussian Mixture (to isolate vessels from background noise). Finaly, we dilated the vessels using a spherical structuring element with a radius of 3mm (6 voxels), and inverted the image to produce a mask excluded vessel-adjacent voxels.

##### Resting-state event detection analyses

To dissociate the influence of DN drive versus dATN drive on spontaneous interactions, we developed an event-triggered analysis of resting-state fMRI data. Because participants were not engaged in any task, resting-state data provide a neutral testbed for evaluating the directionality and spatial specificity of DN–dATN interactions. Building on prior findings that attention during rest naturally alternates between internally and externally oriented states(23, 49, 50), thought to be supported by the DN and dATN respectively, we reasoned that transient “neural events” in these networks could reveal spontaneous DN-driven and dATN-driven communication.

We defined DN-driven events as transient high-activation periods in DN pRFs, and dATN-driven events as high-activation periods in positive-polarity dATN pRFs(98, 99). Event detection was performed independently for each source pRF by taking each pRF’s z-scored resting-state time series and identifying TRs with unusually high activity. Specifically, we considered any time point with a z-value greater than 2.4 (i.e., 99.18 percentile) as a neural event. Results were comparable with a range of thresholds (2.1 < z < 2.9).

At each detected event, we extracted the resting-state time series of the 10 most retinotopically aligned pRFs in the target network—i.e., the 10 with the smallest Euclidean distance in receptive-field center (x, y)—over a 13-TR peri-event window (–6 to +6 TRs). These time series were normalized to the mean of the first four pre-event TRs to allow comparisons across events and conditions. To evaluate the spatial specificity of these interactions, we compared responses in the matched pRFs to those in an “anti-matched” pRFs set: the 10 most distant pRFs in the target network in terms of visual field preference (i.e., pRF center positions). Unlike prior analyses that used random sampling to define unmatched control pRFs, here we used a deterministic anti-match procedure to avoid the computational burden of repeated resampling, given the large number of events.

This procedure was repeated across all relevant source–target network pairs: DN-driven (with dATN as target), dATN-driven (with DN as target), and for both positive and negative DN pRFs. For each event type and match status, we computed the average response in the target pRFs at time zero (event peak). These values were entered into a linear mixed effects model with fixed effects of match status (matched vs. anti-matched) and event direction (DN-driven vs. dATN-driven), and with subject as a random effect. This allowed us to test whether retinotopic alignment modulates DN-dATN interactions during spontaneous events and whether such modulation differs for DN-driven vs. dATN-driven events.

#### Statistical tests

Statistical analyses were implemented in MATLAB (Mathworks, Inc). Given the relatively small number of participants and large number of runs per subject, we used linear mixed effects models to account for both within- and between-subject variability, as described in the relevant sections above. For follow-up tests, we used paired-sample t-tests and corrected for multiple comparisons where appropriate. Corrected p-values are reported where appropriate.

## Competing interests

The authors declare no competing interests.

Preprint: Steel*, Angeli*, Robertson. Retinotopic coding organizes the opponent dynamic between internally and externally oriented brain networks. DOI: 10.1101/2024.09.25.615084.

## Data availability

All data is made publicly available via the Natural Scenes Dataset (https://naturalscenesdataset.org/). All code required to reproduce the results of this paper will be made available in Open Science Framework repository upon publication.

## Acknowledgements

The authors would like to thank the authors of the Natural Scenes Dataset for making these data publicly available. This work was supported by an award from the National Institutes of Mental Health (R01MH130529) to CER. AS was supported by the Neukom Institute for Computational Sciences.

## Author contributions

A.S., P.A., and C.E.R. designed the study, analyzed the data, and interpreted the experiments. A.S. and C.E.R. wrote and edited the manuscript; P.A. edited the manuscript. C.E.R. secured funding.

## Supporting Information

**Fig S1.**
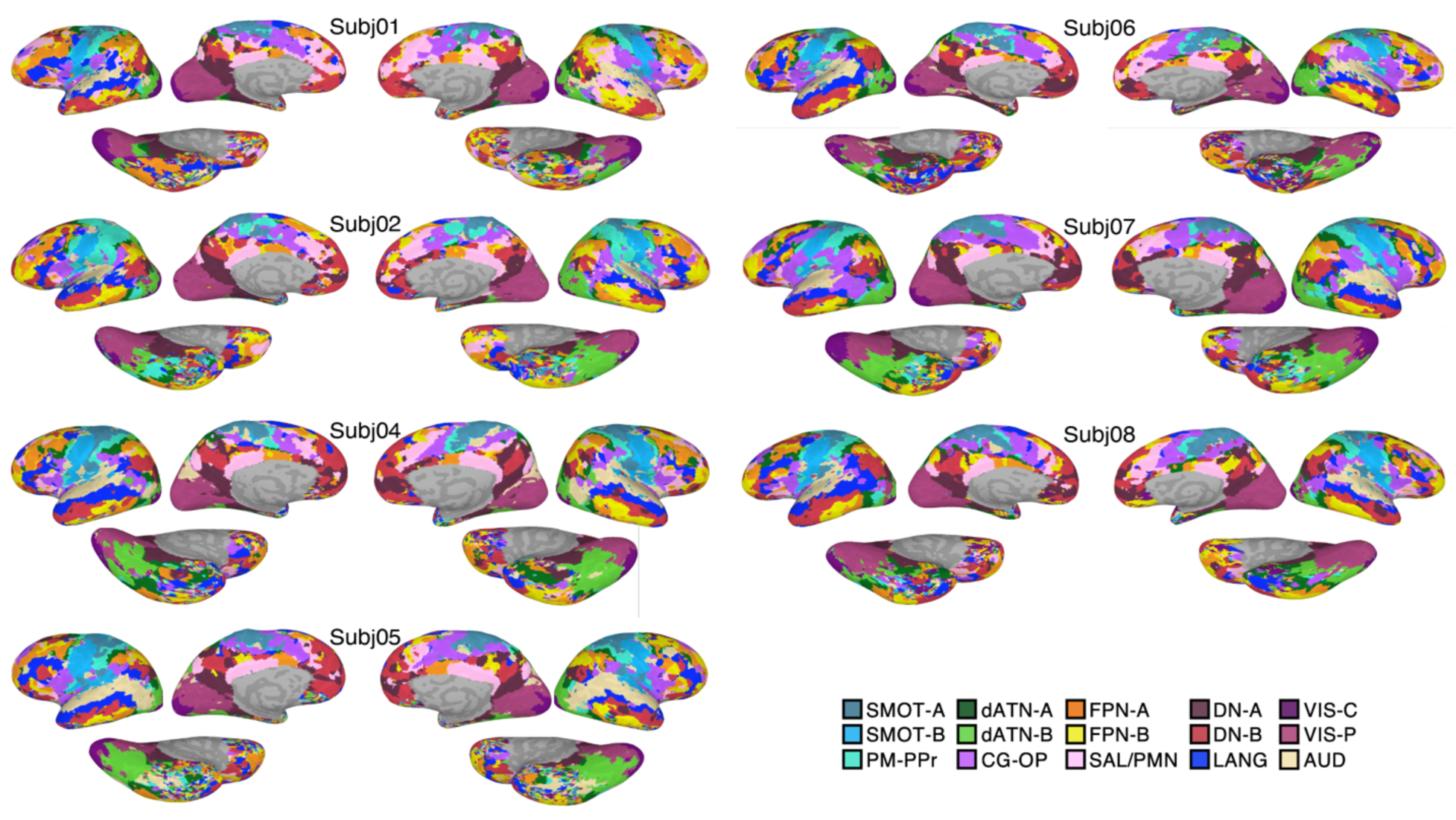
Cortical network parcellations for individual participants. Networks were parcellated based on a 15-network group prior(1, 8) using the multi-session hierarchical Bayesian model (MS-HBM) method(42).

**Fig S2.**
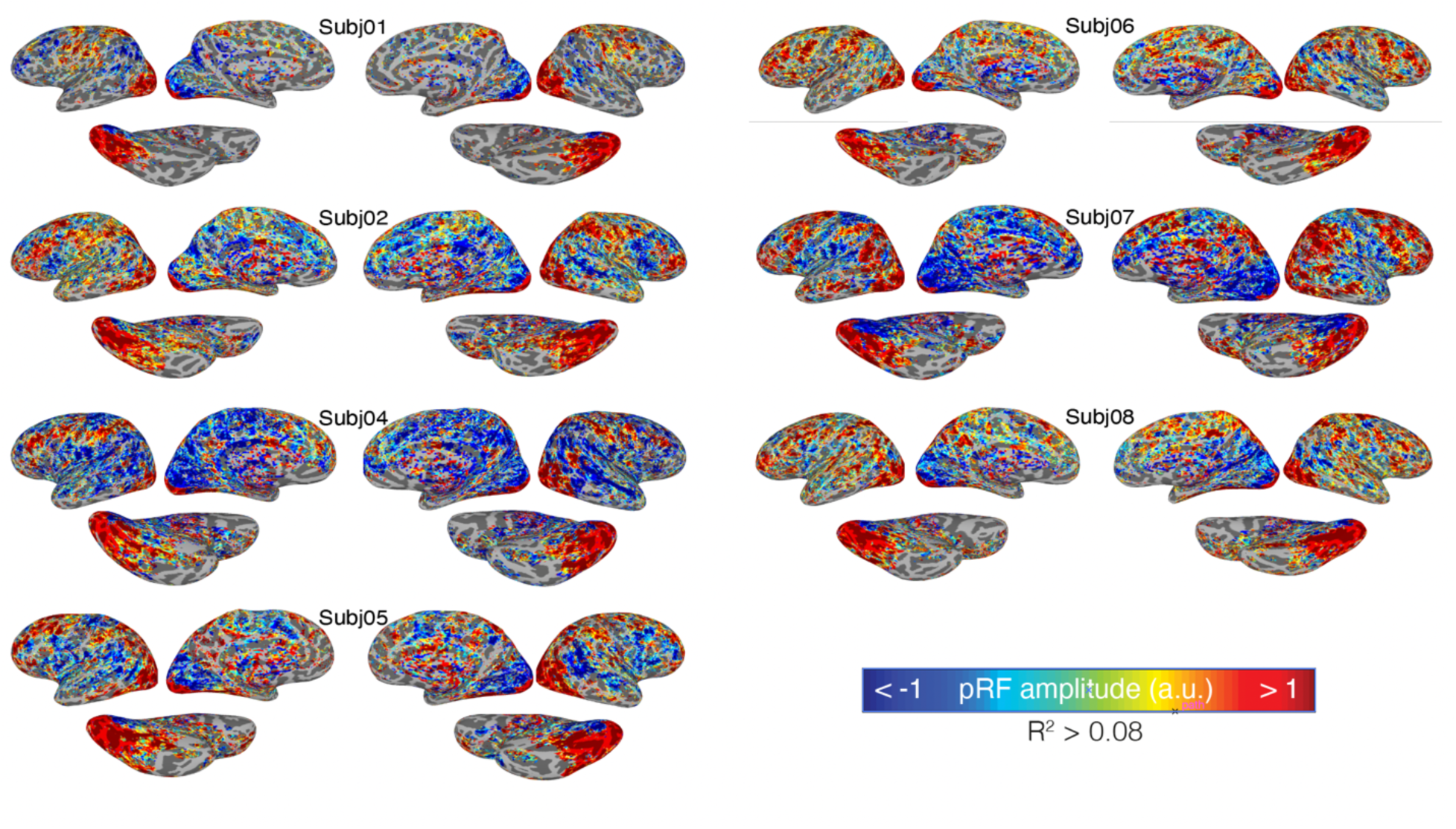
PRF amplitude maps from all participants. Only voxels with greater than 8% variance explained by the pRF model are shown for visualization only.

**Fig. S3.**
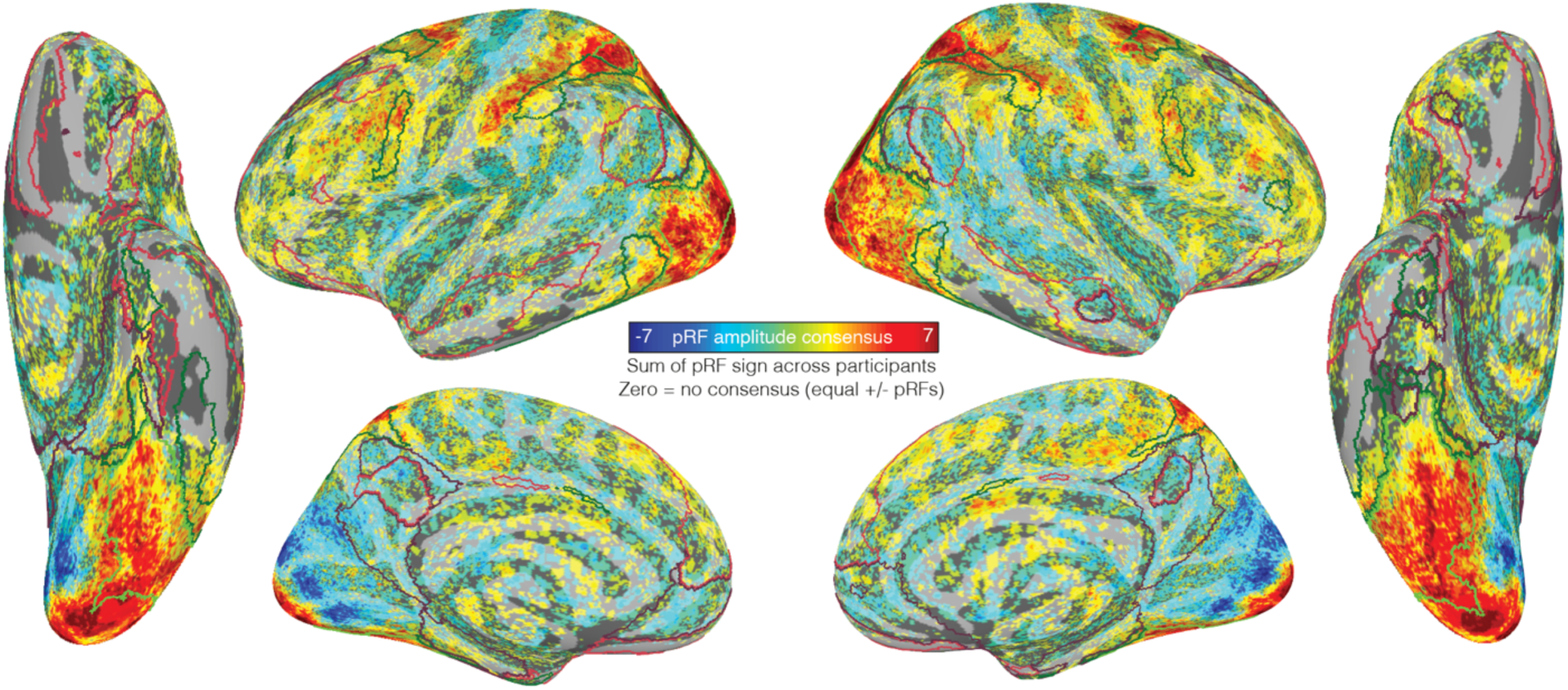
Negative pRFs are clustered in DN-A and DN-B. Brain shows the consensus signed pRF amplitude map, which is the sum of the signed pRF amplitude across participants. Consensus was calculated by binarizing each vertex based on the sign of the visual response: participants with a positive response at a vertex were assigned a +1, participants with a negative response at a vertex were assigned a -1, and participants that did not respond (i.e., had variance explained below our 0.14 threshold) were assigned a 0. We then summed these maps across participants. Vertices with positive values were positive in the majority of participants, vertices with negative values were negative in the majority of participants, and vertices with 0 value showed no consensus. Outlines show DN-A (purple), DN-B (pink), dATN A (light green), and dATN-B (dark green).

**Fig. S4.**
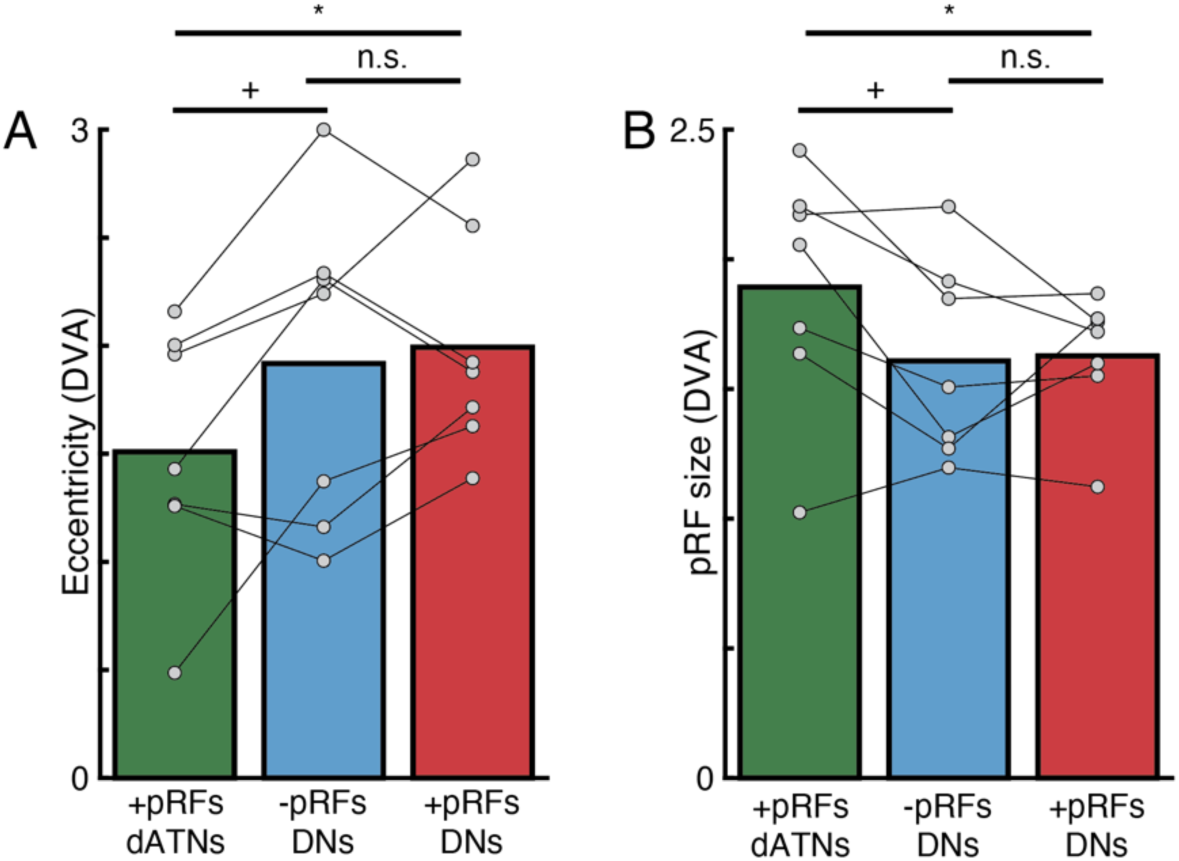
Eccentricity and pRF size differs between DN and dATNs. A. Both positive and negative DN pRFs tend to be more eccentric than dATN pRFs (positive dATN v negative DN pRFs: t(6)=2.27, p=0.06; positive dATN v positive DN pRFs: t(6)=3.04, p=0.023; negative DN v positive DN pRFs: t(6)=0.41, p=0.69). B. Both positive and negative DN pRFs tend to be smaller than dATN pRFs (positive dATN v negative DN pRFs: t(6)=2.36, p=0.056; positive dATN v positive DN pRFs: t(6)=2.46, p=0.049; negative DN v positive DN pRFs: t(6)=0.16, p=0.87) (Fig. S4A). Importantly, these results are averaged across the whole network. Results may differ when individual regions of each network are considered. *=p<0.05, +=p<0.1, n.s. = non-significant

**Fig. S5.**
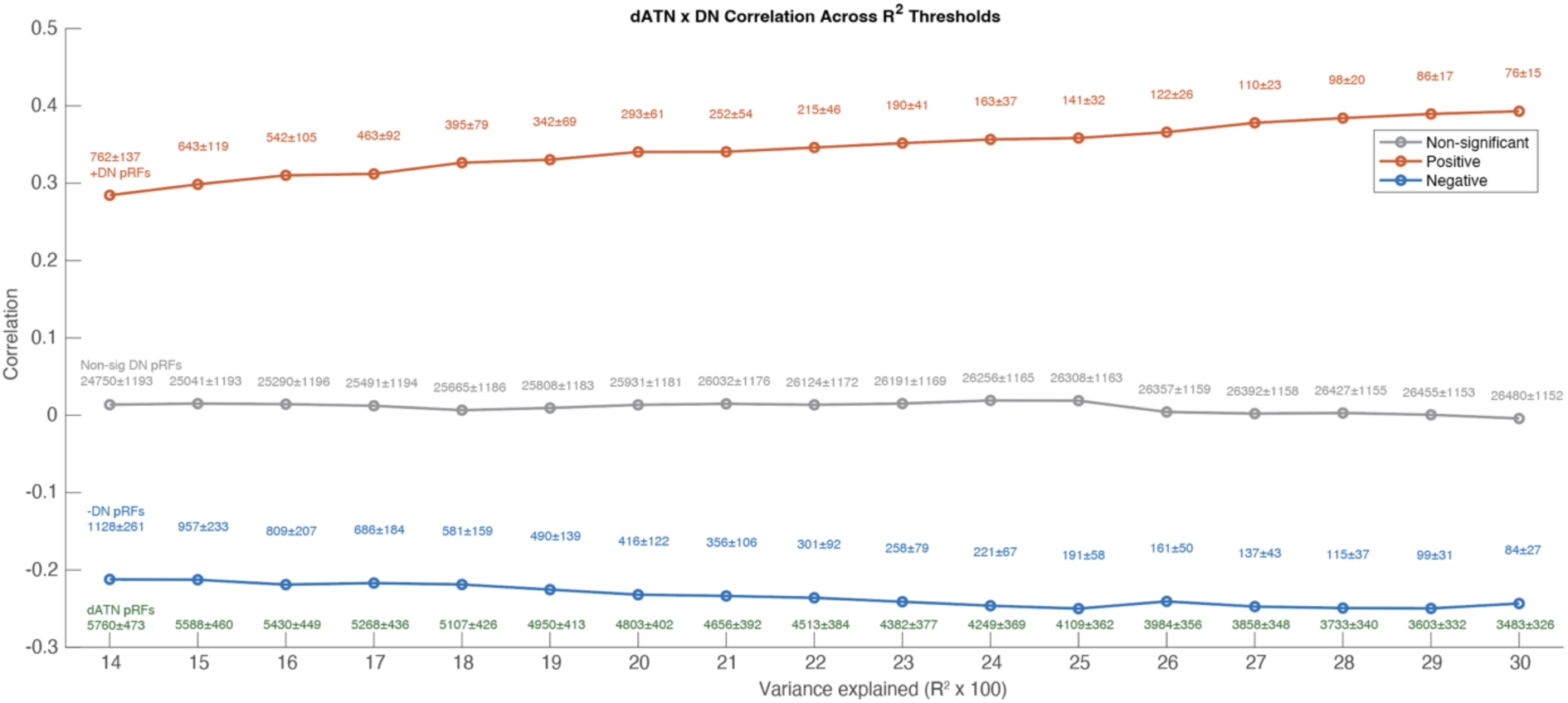
Correlation between dATN and DN voxels across increasingly conservative R^2^ thresholds. Starting from the null-distribution defined threshold of 0.14, increasing R^2^ thresholds results in a more positive correlation between dATN +pRFs and +pRFs in the DN (red), and a more negative correlation with negative DN pRFs (blue). The correlation with non-significant DN voxels remained consistent around zero, mirroring the overall independence of these networks at rest. Text above each point indicates the mean number of DN voxels (with SEM) falling into each significance bin across participants. Red text at bottom of plot indicates the mean number of dATN voxels (with SEM) which passed the threshold and thus was included in the analysis.

**Fig. S6.**
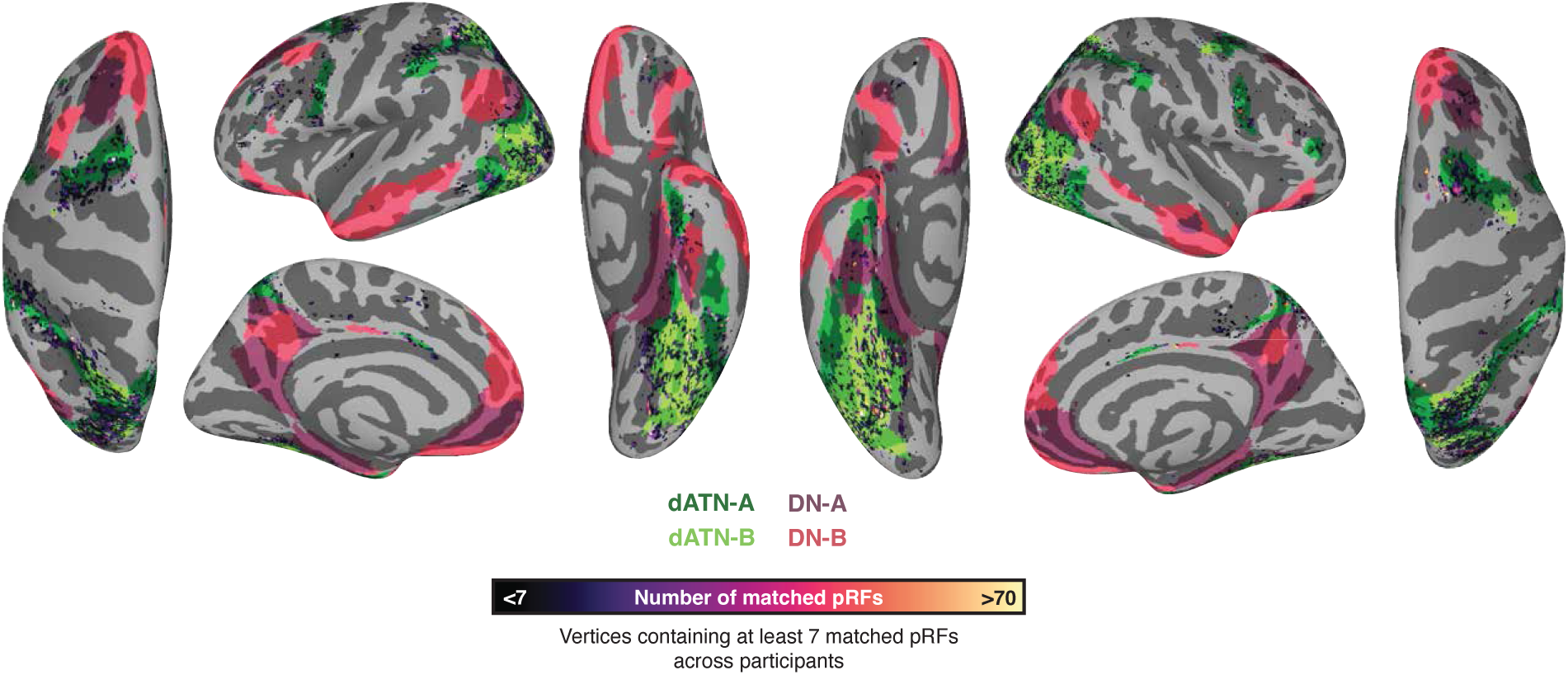
Spatial distribution of positive dATN pRFs matched with negative DN pRFs in cortex across all participants. Vertices are colored based on number of matched pRFs. Only vertices with greater than 7 matched pRFs (i.e., at least one for each subject) are shown. Group prior maps(*1, 8*) dATN A-B and DN A-B are shown for reference. Note that pRFs were matched in individualized dATN of each participant and are not expected to correspond precisely with the group network maps.

**Fig. S7.**
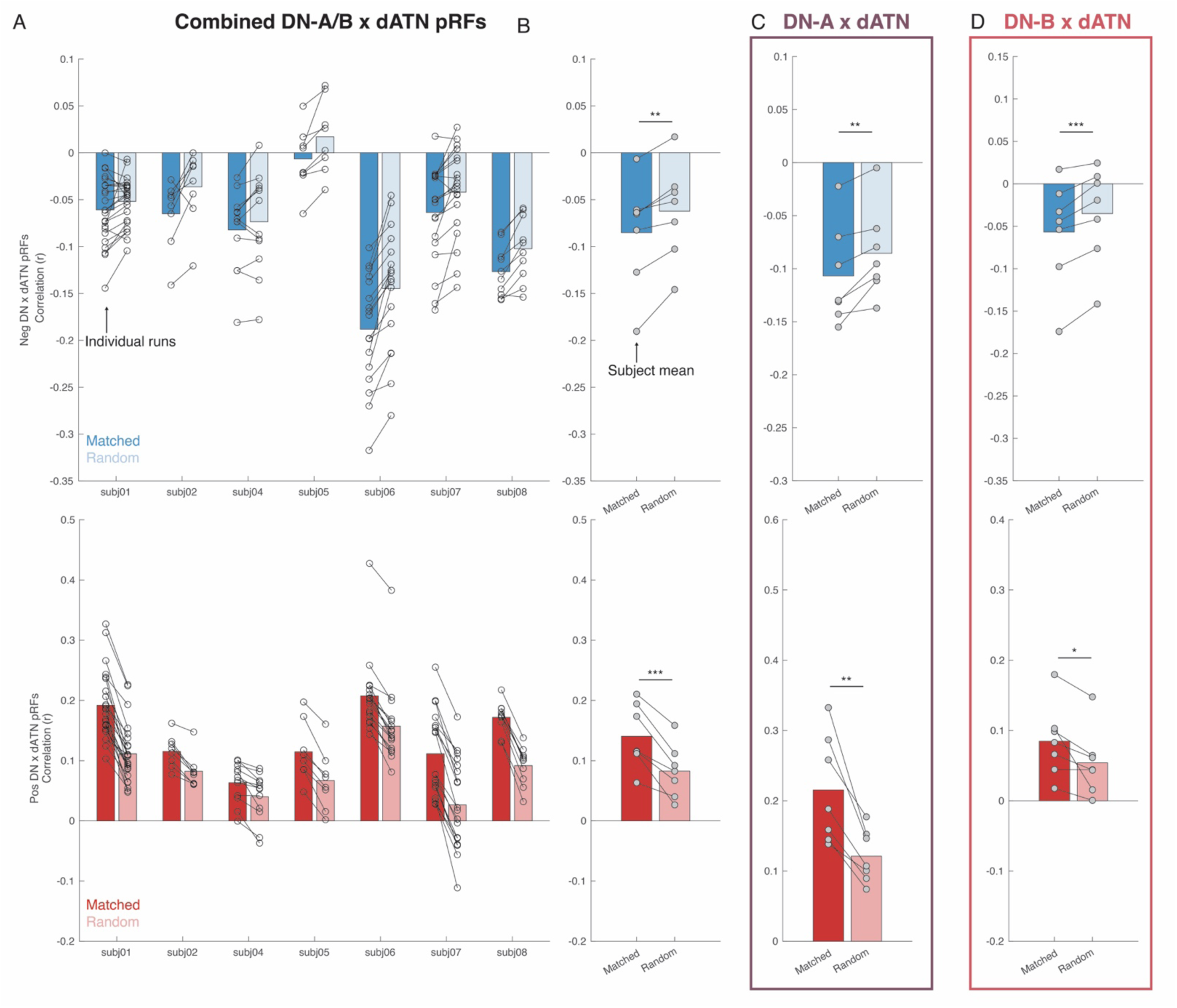
Network pRF matching remains significant after excluding voxels near major veins. Negative (upper) and positive (lower) DN pRFs show retinotopic specific interaction with the dATN. Each set of bars depicts the average correlation of the matched and randomly paired pRFs between the DN and dATN for each participant, with each run shown as connected points. All participants showed greater correlation (either positive or negative) for matched compared with randomly paired pRFs. B. Data shown averaging runs within each participant. For both positive and negative pRFs, the correlation between matched compared with randomly matched pRFs was significantly greater (consistent with their amplitude, ps<0.01). C, D. show the correlation between positive and negative pRFs in the DN with the dATN, split across the sub-networks of the DN-A and B. Both DN-A and B show a significant impact of matching, indicating that both subnetworks interact with the dATN via a retinotopic code (ps<0.05). *: p<0.05, **: p<0.01, ***:p<0.001

**Fig. S8.**
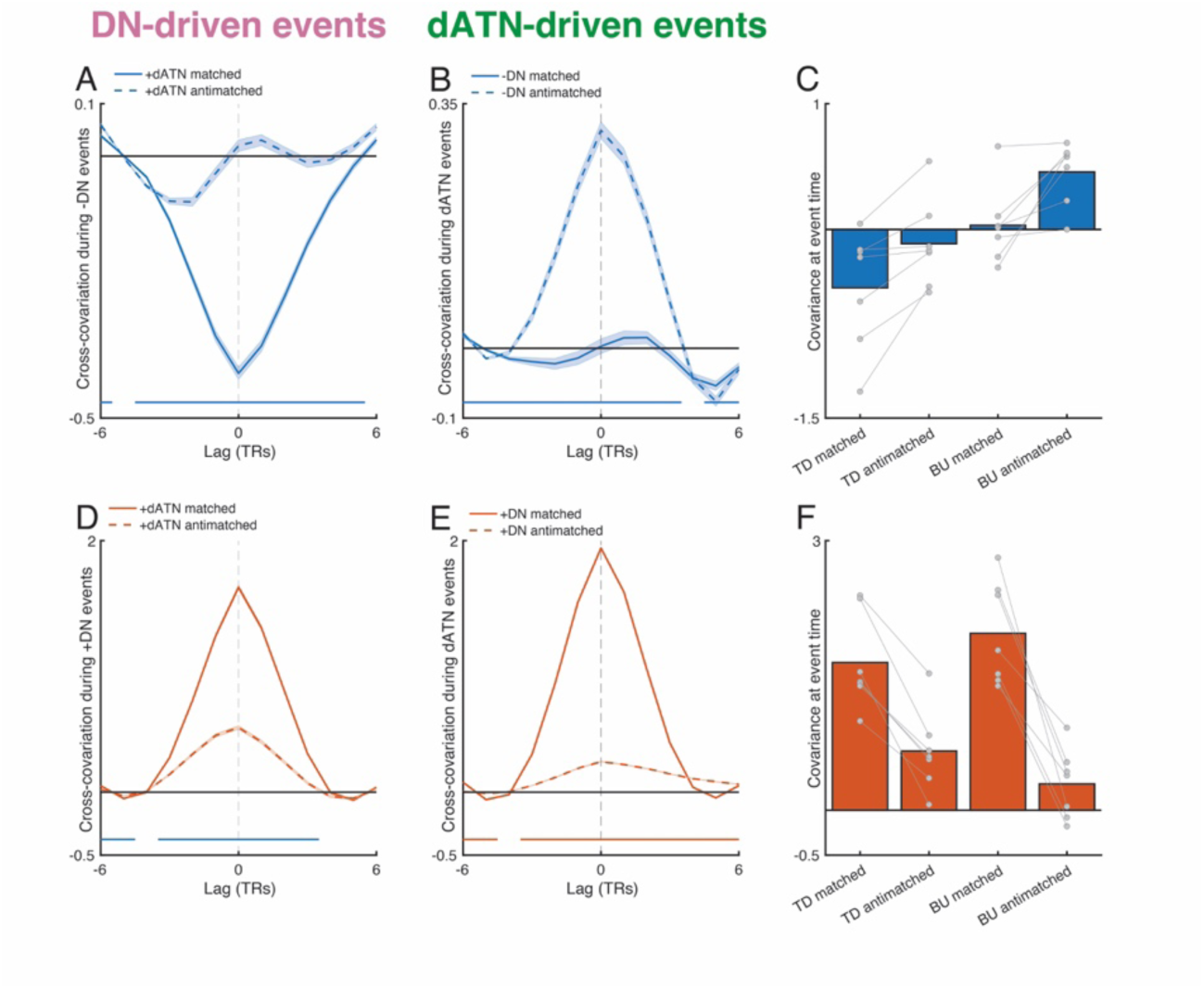
A. Time series show the peri-event cross-covariance of matched and *anti-match*ed dATN pRFs averaged across all participants for events detected in negative DN pRFs (N=78,978 events). B. Time series shows the peri-event cross-covariance in negative DN pRFs during events detected in dATN pRFs (N=541,546). C. Bars show the average cross-covariance at event time of the target areas’ matched and *anti-match*ed pRFs for each participant. Although this difference did not reach significance (F(1,24)=3.94, p=0.058), the results qualitatively support our conclusion – that matched pRFs in the target region have a stronger interaction with the source pRFs. D, E. Evidence for retinotopically-specific excitation of ongoing activity during events DN-driven events detected in positive DN pRFs (D) and dATN-driven events detected in dATN pRFs (E). D. Grand average cross-covariance for matched and *anti-match*ed dATN pRFs during events detected in positive pRFs for all participants. E. Grand average cross-covariance for matched and anitmatched positive DN pRFs during events detected in dATN pRFs. I. For positive DN pRFs paired with dATN pRFs, target area shows retinotopically-specific positive covariance for both DN-driven and dATN-driven events. Bars show the average activation at event time of the target areas’ matched and *anti-match*ed pRFs for each participant. We observed a strong influence of match status on covariance (F(1,25)=25.324, p<0.001; t(6)=7.73, p<0.001). Similar to our findings when examining activation at event time, dATN-driven events had a greater influence from retinotopic coding compared with DN-driven events, evidenced by a greater difference between matched versus *anti-match*ed pRF covariation at event time (F(1,24)=6.22, p=0.02, t(6)=3.99, p=0.007). For time series plots, solid bars along the x-axis indicate periods with a significant difference between matched and *anti-match*ed pRF activity within the target area.

**Fig. S9.**
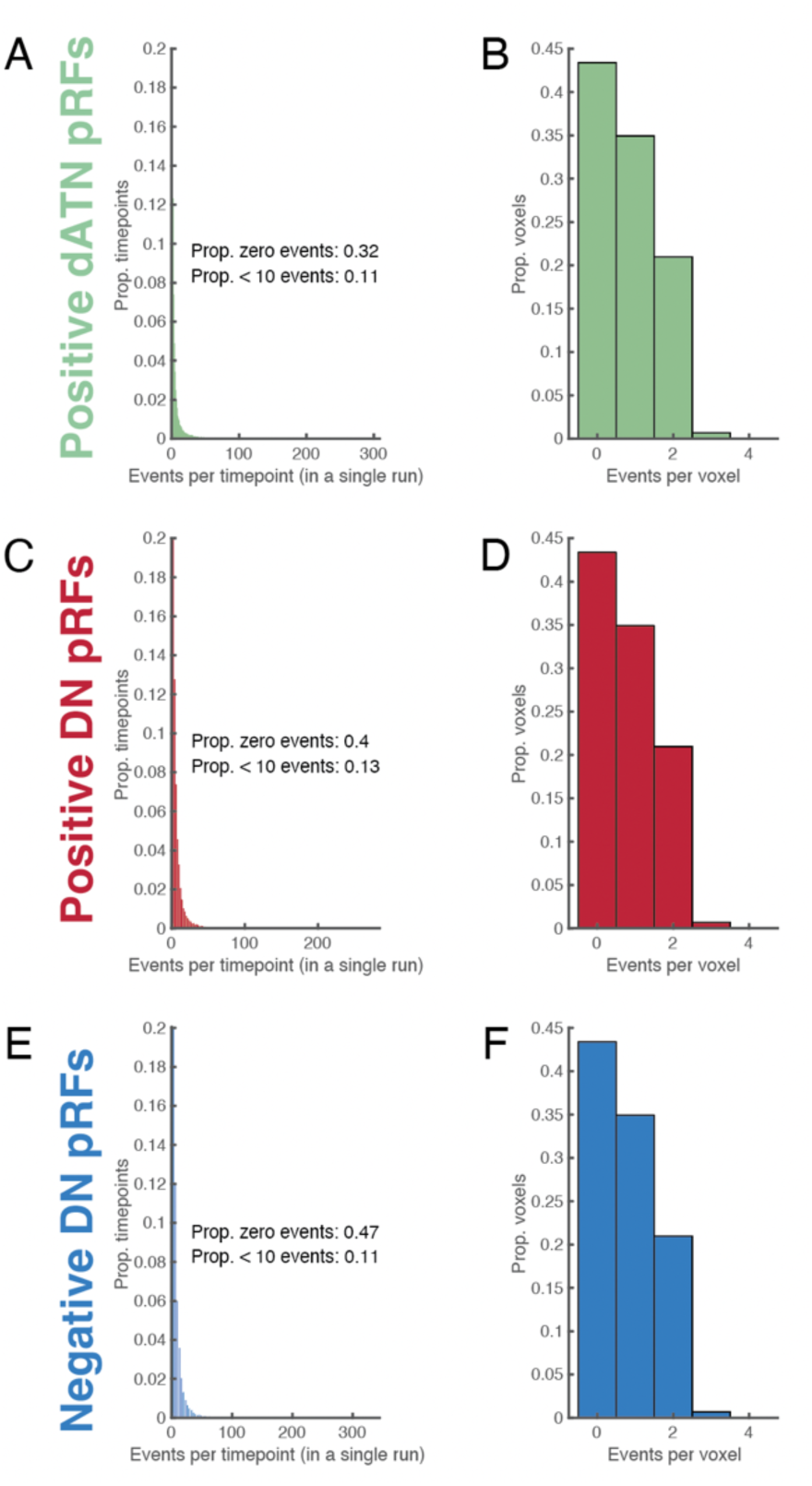
Events are widely distributed in time and across voxels. A,C,E. Number of events per time point combined for top down and bottom up events for dATN, positive DN, and negative DN pRFs. Bars indicate the number of voxel events detected at a given time point within a resting-state run, aggregated across runs. Events were sparsely distributed: most time points contained no events (Proportion = 0.66, bar not shown), with the number of events decreasing approximately exponentially. This is further evidence for a wide temporal distribution of events. B,D,F. Proportion of voxels with events in each run for dATN, positive DN, and negative DN pRFs. No voxel had more than 4 events per run. Data are combined between DN- and and dATN-driven events.

**Fig. S10.**
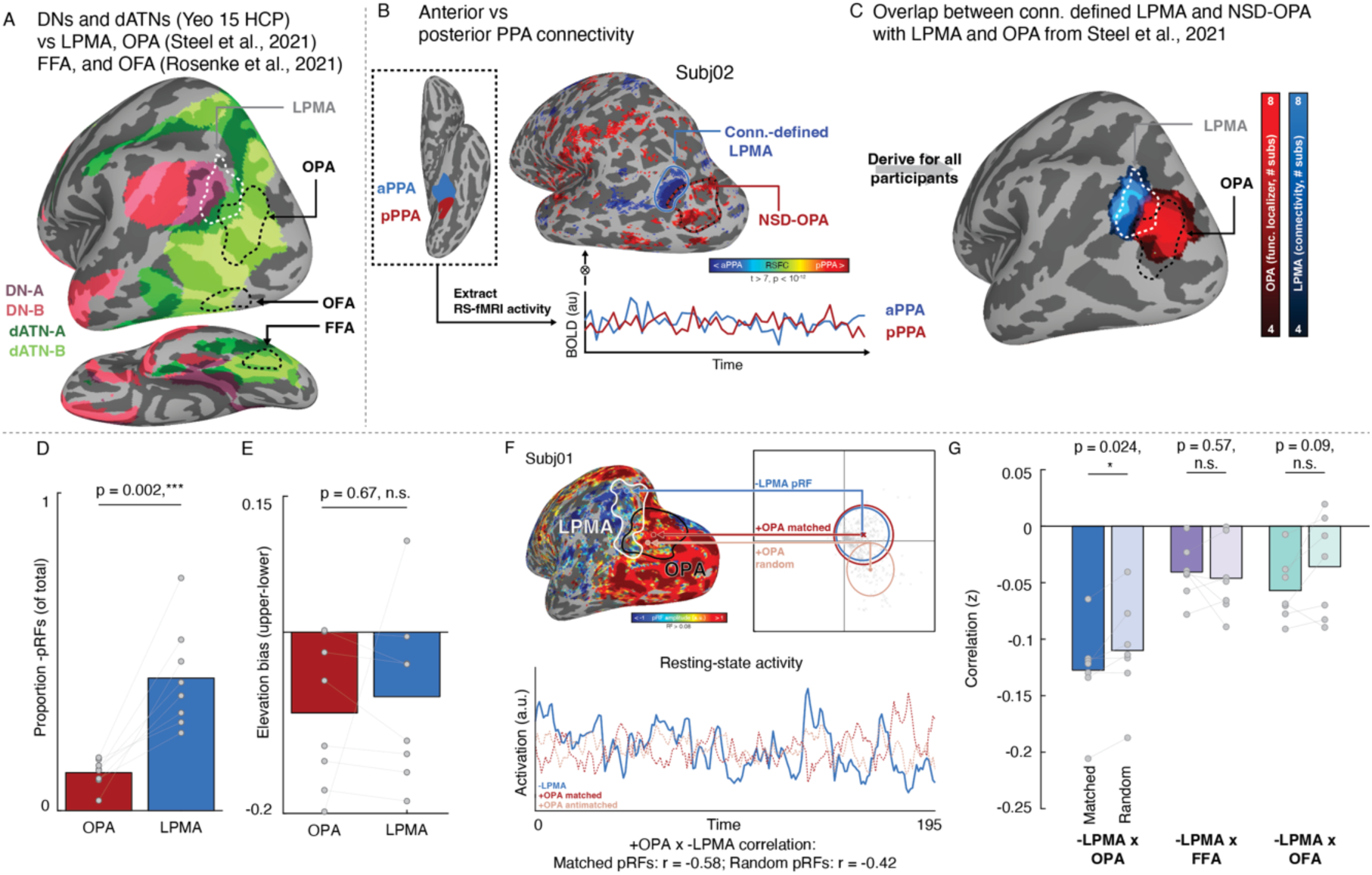
Retinotopic coding structures the spontaneous interaction between functionally-coupled mnemonic and perceptual areas during resting-state fMRI. A. Isolating functionally coupled internally- and externally-oriented brain areas within the DNs and dATNs. We identified brain areas that were specialized in two domains: scenes and faces processing. Specifically, we focused on the lateral place memory area (LPMA; from (36)), white), a memory area in the domain of scene perception at the posterior edge of the DN-A (purple; from (8)). We examined LPMA’s relationship to three different of category-selective visual areas in the dATN (green; from (8)), 1) the occipital place area (OPA; from (36)), an area within the domain of scene perception, along with 2) the occipital face area (OFA) and 3) the fusiform face area (FFA), two areas involved in the domain of face perception (white; from (100)). B-C. We localized LPMA in all participants by contrasting the correlation in resting-state activity between anterior and posterior parahippocampal place area (PPA) (B). This yielded a region in lateral occipital-parietal cortex that overlapped with the LPMA defined in an independent group of participants (C). D-E. Consistent with prior work, the connectivity-defined LPMA had greater concentration of negative pRFs compared to OPA (D), and exhibited a lower visual field bias to OPA (E), consistent with an opponent interaction between these areas during perception. F. We assessed the influence of retinotopic coding on the interaction between negative pRFs in mnemonic and positive pRFs in perceptual areas using the same pRF matching and correlation procedure described above. We compared pRFs within functional domain (scene memory x perception – LPMA to OPA) as well as across domains (scene memory x face perception – LPMA to the occipital face area (OFA) and fusiform face area (FFA)). G. We found that LPMA and OPA were interlocked in a retinotopically-grounded opponent interaction. Resting-state activity of -LPMA and +OPA pRFs was reliably negatively correlated after accounting for variance associated with +LPMA pRFs, and, critically, this negative correlation was stronger for matched compared with randomly matched pRFs (t(6)=3.012, p=0.024). We next tested whether this opponent dynamic was modified by functional domain (i.e., the scene memory area LPMA paired with the face perception areas FFA and OFA), and we observed no significant difference between the distribution of correlation values for matched and randomly-matched pRFs between -LPMA and OFA (t(6)=0.59, p=0.57) and -LPMA and FFA (t(6)=2.01, p=0.09). However, when we compared the difference of matched and randomly-matched pRFs across domains, we found no significant interaction (F(1,19)=0.99,p=0.33).

## Figure S10 Results

Our main results establish that retinotopic coding structures spontaneous interactions between internally- and externally-oriented neural systems in the absence of task demands. However, high-level cortical areas generally associate into networks based on their functional domain. For example, within the visual system, brain areas with differing retinotopic preferences (e.g., the scene-selective areas on the lateral and ventral surfaces of the brain) (43, 101) nevertheless form functional networks based on their apparent preference for specific visual categories (e.g., faces, objects, or scenes)(102, 103). This raises a question: do retinotopic and domain-specific organizational principles interact to facilitate or constrain information flow across internally- and externally-oriented networks? Addressing this question would shed light on mechanisms that enable the brain to integrate information while maintaining functional specialization.

To address this question, we focused on the functional interplay between a set of areas in posterior cerebral cortex that are established models for mnemonic and visual processing in the domains of scene and face perception. Specifically, we considered the mnemonic lateral place memory area (LPMA(36, 78)), an area on the brain’s lateral surface that is implicated in processing mnemonic information relevant to visual scenes at the border between DN-A and the dATNs (Fig 2A). We examined OPA and LPMA because these regions are maximally spatially dissociable (36, 69, 78) and can be localized independently using resting-state data(104, 105). We examined how LPMA activation co-fluctuates with the adjacent, scene-perception area “occipital place area” on the brain’s lateral surface (OPA(106, 107)) compared to two face-perception regions on the lateral and ventral surfaces, the occipital and fusiform face areas (OFA and FFA (108, 109)). At a group level, these perceptual regions are situated within the dATNs and are at the same level of the visual hierarchy (Fig. S10A), but they are differentially associated with the domains of scene (OPA) and face (OFA, FFA) processing, making them ideal model systems to examine the impact of domain-specificity and retinotopic coding in organizing neural activity.

We first defined the mnemonic area LPMA in the NSD participants. To do this, we contrasted functional connectivity between the anterior and posterior halves of the parahippocampal place area using each participant’s resting state data(104, 105) (Fig. S10B), which revealed a cluster in lateral parietal cortex with a similar topographic profile as LPMA based on a group analysis from our prior work (36) (Fig.S8C). Replicating our previous findings(34), this connectivity-defined LPMA had a higher concentration of robust negative pRFs compared to OPA (t(7)=5.26, p=0.002) (Fig. S10D) and exhibited a similar lower visual field bias as OPA (OPA: 8/8 t(7)=3.13, p=0.016; LPMA: 7/8 participants, t(7)=2.11, p=0.07; OPA v LPMA: t(7)=0.441, p=0.67) (Fig. S10E).

We considered whether the retinotopic opponent dynamic we have previously shown in the domain of scenes (i.e., between -LPMA and +OPA pRFs) during perceptual and mnemonic tasks(34) was also present at rest (Fig. S10F). As we observed for the overall negative DN and positive dATN pRFs, we found that -LPMA and +OPA pRFs are interlocked in a retinotopically-grounded opponent interaction. We used the same matching and randomization procedure as we used for the DN.

Remarkably, we found that LPMA and OPA were interlocked in a retinotopically-grounded opponent interaction (Fig. S10G). Resting-state activity of -LPMA and +OPA pRFs was reliably negatively correlated after accounting for variance associated with +LPMA pRFs, and, critically, this negative correlation was stronger for matched compared with randomly matched pRFs (t(6)=3.012, p=0.024). We next tested whether this opponent dynamic was modified by functional domain (i.e., the scene memory area LPMA paired with the face perception areas FFA and OFA), and we observed no significant difference between the distribution of correlation values for matched and randomly-matched pRFs between -LPMA and OFA (t(6)=0.59, p=0.57) and -LPMA and FFA (t(6)=2.01, p=0.09). However, when we compared the difference of matched and randomly-matched pRFs across domains, we found no significant interaction (F(1,19)=0.99,p=0.33).

One possibility is that the quality of the retinotopic matching between pRFs in different areas might contribute to the within versus across domain distinction in retinotopic coding. We checked whether there was a difference in the match quality between the top 10 best matched pRFs between connectivity-defined LPMA and OPA compared with the face perception areas FFA and OFA. For this analysis, we computed the euclidean distance between all pRFs centers between regions, and we took the median of the best pRF center matches. For example, if a participant had 162 negative pRFs LPMA and 90 pRFs in OPA, we found the top 10 best matched OPA pRFs for each LPMA pRF, and then took the median distance of these 1620 matches. This gave us the most representative distance for matched pRFs for each participant.

Overall, we found that the quality of pRF matching was better for within domain (LPMA to OPA) compared with across domains (LPMA to OFA and FFA) (Negative pRFs: t(6)=5.86, p=0.001; positive pRFs: t(6)=6.79, p<0.001). This suggests that the quality of the matches does depend on the stimulus domain. This fits with the notion that visual response properties are inherited by upstream areas, and that these response properties depend in part on the nature of the stimuli within a domain.

These results suggest that retinotopic coding structures the interaction between brain areas if they share a functional domain, but more studies are needed to confirm whether this is specific to within-domain matching.

## Figure S10 Methods

### Defining functional regions of interest (PPA, OPA, OFA, FFA)

We used the volumetric functional regions of interest provided with the NSD at 1.8mm resolution. Specifically, we used the parahippocampal place area (PPA), occipital place area (OPA), iog-faces (referred to here as occipital face area (OFA)), and pfus-faces (fusiform face area, FFA1) regions of interest. In brief, these regions were defined in the NSD using within-subject data collected from 6 runs of a multi-category visual localizer paradigm (37).

### Defining functional region LPMA

Because the NSD did not include mnemonic localizers (36), we used the resting-state data to define the lateral place memory area (LPMA). Briefly, the LPMA is a region anterior to OPA and near caudal inferior parietal lobule(104, 105) that selectively responds during recall of personally familiar places compared to other stimulus types. We have previous shown that OPA and LPMA are functionally-linked and work jointly to process knowledge of visuospatial context out of view during scene perception.

Prior work has suggested that a mnemonic area linked to scenes on the lateral surface can be localized by comparing resting-state co-fluctuations of anterior versus posterior PPA (aPPA and pPPA, respectively)(19, 104, 105), and we adopted that approach here. We preprocessed the resting-state fixation fMRI data, runs 1 and 14 from NSD sessions 22 and 23, in all participants (prior to data exclusion) and extracted the average time series of aPPA, pPPA, aFFA, and pFFA. We used these time series as regressors in a general linear model, and we compared the beta-values from the aPPA and pPPA. We considered any voxels with a t-statistic > 5 within posterior parietal-occipital cortex on the lateral surface as an LPMA ROI (Individual ROIs can be found in Supplemental Fig. 4). Across subjects, there was considerable overlap between this connectivity-defined area and a group-level LPMA defined based on our prior work (Fig. 2).

## References

1. B. T. Thomas Yeo, et al., The organization of the human cerebral cortex estimated by intrinsic functional connectivity. J. Neurophysiol. 106, 1125–1165 (2011).

2. D. S. Margulies, et al., Situating the default-mode network along a principal gradient of macroscale cortical organization. Proc. Natl. Acad. Sci. U. S. A. 113, 12574–12579 (2016).

3. R. M. Braga, R. L. Buckner, Parallel Interdigitated Distributed Networks within the Individual Estimated by Intrinsic Functional Connectivity. Neuron 95, 457–471.e5 (2017).

4. E. M. Gordon, et al., Precision Functional Mapping of Individual Human Brains. Neuron 95, 791–807.e7 (2017).

5. C. Ranganath, M. Ritchey, Two cortical systems for memory-guided behaviour. Nat. Rev. Neurosci. [Preprint] (2012). [Accessed 11 March 2020].

6. J. R. Andrews-Hanna, J. S. Reidler, J. Sepulcre, R. Poulin, R. L. Buckner, Functional-Anatomic Fractionation of the Brain’s Default Network. Neuron 65, 550–562 (2010).

7. A. J. Barnett, et al., Intrinsic connectivity reveals functionally distinct cortico-hippocampal networks in the human brain. PLoS Biol. 19, e3001275 (2021).

8. J. Du, et al., Organization of the human cerebral cortex estimated within individuals: networks, global topography, and function. J. Neurophysiol. 131, 1014–1082 (2024).

9. T. O. Laumann, et al., Functional System and Areal Organization of a Highly Sampled Individual Human Brain. Neuron 87, 657–670 (2015).

10. M. M. Chun, J. D. Golomb, N. B. Turk-Browne, A Taxonomy of External and Internal Attention. Annu. Rev. Psychol. 62, 73–101 (2011).

11. M. L. Dixon, et al., Interactions between the default network and dorsal attention network vary across default subsystems, time, and cognitive states. Neuroimage 147, 632–649 (2017).

12. M. D. Fox, et al., The human brain is intrinsically organized into dynamic, anticorrelated functional networks. Proc. Natl. Acad. Sci. U. S. A. 102, 9673–9678 (2005).

13. M. E. Raichle, The Brain’s Default Mode Network. Annu. Rev. Neurosci. 38, 433–447 (2015).

14. M. D. Fox, M. Corbetta, A. Z. Snyder, J. L. Vincent, M. E. Raichle, Spontaneous neuronal activity distinguishes human dorsal and ventral attention systems. Proceedings of the National Academy of Sciences 103, 10046–10051 (2006).

15. M. E. Raichle, et al., A default mode of brain function. Proceedings of the National Academy of Sciences 98, 676–682 (2001).

16. E. Fedorenko, J. Duncan, N. Kanwisher, Broad domain generality in focal regions of frontal and parietal cortex. Proceedings of the National Academy of Sciences 110, 16616–16621 (2013).

17. S. Vossel, J. J. Geng, G. R. Fink, Dorsal and Ventral Attention Systems: Distinct Neural Circuits but Collaborative Roles. The Neuroscientist 20, 150–159 (2013).

18. G. L. Shulman, et al., Common Blood Flow Changes across Visual Tasks: II. Decreases in Cerebral Cortex. J. Cogn. Neurosci. 9, 648–663 (1997).

19. E. H. Silson, A. Steel, A. Kidder, A. W. Gilmore, C. I. Baker, Distinct subdivisions of human medial parietal cortex support recollection of people and places. Elife 8 (2019).

20. L. M. DiNicola, R. M. Braga, R. L. Buckner, Parallel distributed networks dissociate episodic and social functions within the individual. J. Neurophysiol. 123, 1144–1179 (2020).

21. P. P. Thakral, K. P. Madore, D. L. Schacter, A Role for the Left Angular Gyrus in Episodic Simulation and Memory. J. Neurosci. 37, 8142–8149 (2017).

22. A. W. Gilmore, et al., Dynamic Content Reactivation Supports Naturalistic Autobiographical Recall in Humans. Journal of Neuroscience 41, 153–166 (2021).

23. K. Christoff, A. M. Gordon, J. Smallwood, R. Smith, J. W. Schooler, Experience sampling during fMRI reveals default network and executive system contributions to mind wandering. Proc. Natl. Acad. Sci. U. S. A. 106, 8719–8724 (2009).

24. M. D. Greicius, B. Krasnow, A. L. Reiss, V. Menon, Functional connectivity in the resting brain: A network analysis of the default mode hypothesis. Proceedings of the National Academy of Sciences 100, 253–258 (2003).

25. V. Menon, 20 years of the default mode network: A review and synthesis. Neuron 111, 2469–2487 (2023).

26. Z. S. Saad, et al., Trouble at Rest: How Correlation Patterns and Group Differences Become Distorted After Global Signal Regression. Brain Connect. 2, 25–32 (2012).

27. C. Murphy, et al., Modes of operation: A topographic neural gradient supporting stimulus dependent and independent cognition. Neuroimage 186, 487–496 (2019).

28. C. Murphy, et al., Distant from input: Evidence of regions within the default mode network supporting perceptually-decoupled and conceptually-guided cognition. Neuroimage 171, 393–401 (2018).

29. J. L. S. Bellmund, P. Gärdenfors, E. I. Moser, C. F. Doeller, Navigating cognition: Spatial codes for human thinking. Science (1979). 362, eaat6766 (2018).

30. S. F. Popham, et al., Visual and linguistic semantic representations are aligned at the border of human visual cortex. Nature Neuroscience *2021 24:11* 24, 1628–1636 (2021).

31. M. Szinte, T. Knapen, Visual Organization of the Default Network. Cerebral Cortex 30, 3518–3527 (2020).

32. T. Knapen, Topographic connectivity reveals task-dependent retinotopic processing throughout the human brain. Proc. Natl. Acad. Sci. U. S. A. 118 (2021).

33. P. Christiaan Klink, X. Chen, W. Vanduffel, P. R. Roelfsema, Population receptive fields in non-human primates from whole-brain fmri and large-scale neurophysiology in visual cortex. Elife 10 (2021).

34. A. Steel, E. H. Silson, B. D. Garcia, C. E. Robertson, A retinotopic code structures the interaction between perception and memory systems. Nat. Neurosci. 27, 339–347 (2024).

35. I. I. A. Groen, T. M. Dekker, T. Knapen, E. H. Silson, Visuospatial coding as ubiquitous scaffolding for human cognition. Trends Cogn. Sci. 26, 81–96 (2022).

36. A. Steel, M. M. Billings, E. H. Silson, C. E. Robertson, A network linking scene perception and spatial memory systems in posterior cerebral cortex. Nature Communications 2021 *12:1* 12, 1–13 (2021).

37. E. J. Allen, et al., A massive 7T fMRI dataset to bridge cognitive neuroscience and artificial intelligence. Nature Neuroscience *2021 25:1* 25, 116–126 (2021).

38. S. O. Dumoulin, B. A. Wandell, Population receptive field estimates in human visual cortex. Neuroimage 39, 647 (2008).

39. R. Kong, et al., Individual-Specific Areal-Level Parcellations Improve Functional Connectivity Prediction of Behavior. Cerebral Cortex 31, 4477–4500 (2021).

40. L. Griffanti, et al., ICA-based artefact removal and accelerated fMRI acquisition for improved resting state network imaging. Neuroimage 95, 232–247 (2014).

41. L. Griffanti, et al., Hand classification of fMRI ICA noise components. Neuroimage 154, 188–205 (2017).

42. R. Kong, et al., Spatial Topography of Individual-Specific Cortical Networks Predicts Human Cognition, Personality, and Emotion. Cerebral Cortex 29, 2533–2551 (2019).

43. E. H. Silson, A. W. Y. Chan, R. C. Reynolds, D. J. Kravitz, C. I. Baker, A retinotopic basis for the division of high-level scene processing between lateral and ventral human occipitotemporal cortex. Journal of Neuroscience 35, 11921–11935 (2015).

44. S. M. Smith, et al., Functional connectomics from resting-state fMRI. Trends Cogn. Sci. 17, 666–682 (2013).

45. L. M. DiNicola, R. L. Buckner, Precision estimates of parallel distributed association networks: evidence for domain specialization and implications for evolution and development. Curr. Opin. Behav. Sci. 40, 120–129 (2021).

46. L. M. DiNicola, R. M. Braga, R. L. Buckner, Parallel Distributed Networks Dissociate Episodic and Social Functions Within the Individual. bioRxiv 733048 (2019). 10.1101/733048.

47. L. M. DiNicola, O. I. Ariyo, R. L. Buckner, Functional specialization of parallel distributed networks revealed by analysis of trial-to-trial variation in processing demands. J. Neurophysiol. 129, 17–40 (2023).

48. C. L. Scrivener, E. H. Silson, Opponent visuospatial coding structures responses during memory recall and visual perception in medial parietal cortex. Imaging Neuroscience 3 (2025).

49. J. Gonzalez-Castillo, J. W. Y. Kam, C. W. Hoy, P. A. Bandettini, How to Interpret Resting-State fMRI: Ask Your Participants. The Journal of Neuroscience 41, 1130–1141 (2021).

50. L. Van Calster, A. D’Argembeau, E. Salmon, F. Peters, S. Majerus, Fluctuations of Attentional Networks and Default Mode Network during the Resting State Reflect Variations in Cognitive States: Evidence from a Novel Resting-state Experience Sampling Method. J. Cogn. Neurosci. 29, 95–113 (2017).

51. S. E. Favila, B. A. Kuhl, J. Winawer, Perception and memory have distinct spatial tuning properties in human visual cortex. Nature Communications *2022 13:1* 13, 1–21 (2022).

52. J. L. Breedlove, G. St-Yves, C. A. Olman, T. Naselaris, Generative Feedback Explains Distinct Brain Activity Codes for Seen and Mental Images. Current Biology 1–14 (2020). 10.1016/j.cub.2020.04.014.

53. T. J. Buschman, M. Siegel, J. E. Roy, E. K. Miller, Neural substrates of cognitive capacity limitations. Proc. Natl. Acad. Sci. U. S. A. 108, 11252–11255 (2011).

54. A. Libby, T. J. Buschman, Rotational dynamics reduce interference between sensory and memory representations. Nature Neuroscience *2021 24:5* 24, 715–726 (2021).

55. J. D. Semedo, A. Zandvakili, C. K. Machens, B. M. Yu, A. Kohn, Cortical areas interact through a communication subspace. Neuron 102, 249 (2019).

56. G. G. Gregoriou, S. J. Gotts, H. Zhou, R. Desimone, High-frequency, long-range coupling between prefrontal and visual cortex during attention. Science 324, 1207–1210 (2009).

57. C. A. Bosman, et al., Attentional stimulus selection through selective synchronization between monkey visual areas. Neuron 75, 875–888 (2012).

58. N. Hedger, T. Naselaris, K. Kay, T. Knapen, Vicarious body maps bridge vision and touch in the human brain. Nature 650, 173–181 (2026).

59. D. S. Margulies, et al., Situating the default-mode network along a principal gradient of macroscale cortical organization. Proc. Natl. Acad. Sci. U. S. A. 113, 12574–12579 (2016).

60. R. L. Buckner, L. M. DiNicola, The brain’s default network: updated anatomy, physiology and evolving insights. Nat. Rev. Neurosci. (2019). 10.1038/s41583-019-0212-7.

61. Y. Yeshurun, M. Nguyen, U. Hasson, The default mode network: where the idiosyncratic self meets the shared social world. Nature Reviews Neuroscience *2021 22:3* 22, 181–192 (2021).

62. R. N. Spreng, W. D. Stevens, J. P. Chamberlain, A. W. Gilmore, D. L. Schacter, Default network activity, coupled with the frontoparietal control network, supports goal-directed cognition. Neuroimage 53, 303–317 (2010).

63. R. N. Spreng, R. A. Mar, A. S. N. Kim, The common neural basis of autobiographical memory, prospection, navigation, theory of mind, and the default mode: A quantitative meta-analysis. J. Cogn. Neurosci. 21, 489–510 (2009).

64. B. L. Foster, et al., A tripartite view of the posterior cingulate cortex. Nat. Rev. Neurosci. 24, 173–189 (2023).

65. B. A. Wandell, S. O. Dumoulin, A. A. Brewer, Visual field maps in human cortex. Neuron 56, 366–383 (2007).

66. J. S. Taube, The Head Direction Signal: Origins and Sensory-Motor Integration. Annu. Rev. Neurosci. 30, 181–207 (2007).

67. A. Finkelstein, et al., Three-dimensional head-direction coding in the bat brain. Nature 517, 159–164 (2015).

68. M. Nau, T. Navarro Schröder, M. Frey, C. F. Doeller, Behavior-dependent directional tuning in the human visual-navigation network. Nat. Commun. 11, 3247 (2020).

69. A. Steel, D. Prasad, B. D. Garcia, C. E. Robertson, Relating scene memory and perception activity to functional properties, networks, and landmarks of posterior cerebral cortex - a probabilistic atlas. The Journal of Neuroscience e0028252025 (2025). 10.1523/JNEUROSCI.0028-25.2025.

70. A. M. Bastos, et al., Canonical Microcircuits for Predictive Coding. Neuron 76, 695–711 (2012).

71. T. D. Albright, On the perception of probable things: neural substrates of associative memory, imagery, and perception. Neuron 74, 227–245 (2012).

72. H. C. Barron, T. P. Vogels, T. E. Behrens, M. Ramaswami, E. D. by Thomas Albright, Inhibitory engrams in perception and memory. 114, 6666–6674 (2017).

73. J. M. Huntenburg, P. L. Bazin, D. S. Margulies, Large-Scale Gradients in Human Cortical Organization. Trends Cogn. Sci. 22, 21–31 (2018).

74. M. Zhang, et al., Sender–receiver subdivisions of the default mode network in perceptual and memory-guided cognition. Proceedings of the National Academy of Sciences 123 (2026).

75. Y. Wu, E. Podvalny, M. Levinson, B. J. He, Network mechanisms of ongoing brain activity’s influence on conscious visual perception. Nat. Commun. 15, 5720 (2024).

76. T.-A. E. Nghiem, et al., Space wandering in the rodent default mode network. Proceedings of the National Academy of Sciences 121 (2024).

77. A. Steel, C. Thomas, A. Trefler, G. Chen, C. I. Baker, Finding the baby in the bath water – evidence for task-specific changes in resting state functional connectivity evoked by training. Neuroimage 188, 524–538 (2019).

78. A. Steel, B. D. Garcia, K. Goyal, A. Mynick, C. E. Robertson, Scene Perception and Visuospatial Memory Converge at the Anterior Edge of Visually Responsive Cortex. The Journal of Neuroscience 43, 5723 (2023).

79. T. Çukur, S. Nishimoto, A. G. Huth, J. L. Gallant, Attention during natural vision warps semantic representation across the human brain. Nat. Neurosci. 16, 763–770 (2013).

80. M. Arcaro, M. Livingstone, A Whole-Brain Topographic Ontology. Annu. Rev. Neurosci. 47, 21–40 (2024).

81. M. Peer, I. K. Brunec, N. S. Newcombe, R. A. Epstein, Structuring Knowledge with Cognitive Maps and Cognitive Graphs. Trends Cogn. Sci. (2020). 10.1016/j.tics.2020.10.004.

82. A. G. Huth, W. A. De Heer, T. L. Griffiths, F. E. Theunissen, J. L. Gallant, Natural speech reveals the semantic maps that tile human cerebral cortex. Nature 532, 453–458 (2016).

83. N. Hedger, T. Naselaris, K. Kay, T. Knapen, Vicarious Somatotopic Maps Tile Visual Cortex. bioRxiv 2024.10.21.619382 (2024). 10.1101/2024.10.21.619382.

84. C. A. Pizka, et al., Population receptive field properties of the human visual claustrum zone. [Preprint] (2026).

85. D. M. van Es, W. van der Zwaag, T. Knapen, Topographic Maps of Visual Space in the Human Cerebellum. Current Biology 29, 1689–1694.e3 (2019).

86. P. A. Angeli, A. Steel, E. H. Silson, C. E. Robertson, Positive and Negative Retinotopic Codes in the Human Hippocampus. bioRxiv 2024.09.27.615397 (2024). 10.1101/2024.09.27.615397.

87. K. N. Kay, J. Winawer, A. Mezer, B. A. Wandell, Compressive spatial summation in human visual cortex. J. Neurophysiol. 110, 481–494 (2013).

88. W. Zuiderbaan, B. M. Harvey, S. O. Dumoulin, Modeling center-surround configurations in population receptive fields using fMRI. J. Vis. 12, 10–10 (2012).

89. A. M. Dale, B. Fischl, M. I. Sereno, Cortical surface-based analysis: I. Segmentation and surface reconstruction. Neuroimage 9, 179–194 (1999).

90. B. Fischl, et al., Whole brain segmentation: Automated labeling of neuroanatomical structures in the human brain. Neuron 33, 341–355 (2002).

91. Z. S. Saad, R. C. Reynolds, SUMA. Neuroimage 62, 768–773 (2012).

92. R. W. Cox, AFNI: Software for analysis and visualization of functional magnetic resonance neuroimages. Computers and Biomedical Research 29, 162–173 (1996).

93. P. A. Taylor, et al., A Set of FMRI Quality Control Tools in AFNI: Systematic, in-depth and interactive QC with afni_proc.py and more. bioRxiv 2024.03.27.586976 (2024). 10.1101/2024.03.27.586976.

94. A. B. Beckers, et al., Comparing the efficacy of data-driven denoising methods for a multi-echo fMRI acquisition at 7T. Neuroimage 280, 120361 (2023).

95. M. Jenkinson, C. F. Beckmann, T. E. J. Behrens, M. W. Woolrich, S. M. Smith, FSL. Neuroimage 62, 782–790 (2012).

96. S. M. Smith, et al., Advances in functional and structural MR image analysis and implementation as FSL in NeuroImage, (Academic Press, 2004), pp. S208–S219.

97. I. Alvarez, B. de Haas, C. A. Clark, G. Rees, D. Samuel Schwarzkopf, Comparing different stimulus configurations for population receptive field mapping in human fMRI. Front. Hum. Neurosci. 9, 121251 (2015).

98. E. M. Gordon, et al., A somato-cognitive action network alternates with effector regions in motor cortex. Nature 617, 351–359 (2023).

99. A. Mitra, A. Z. Snyder, T. Blazey, M. E. Raichle, Lag threads organize the brain’s intrinsic activity. Proceedings of the National Academy of Sciences 112, E2235–E2244 (2015).

100. M. Rosenke, R. van Hoof, J. van den Hurk, K. Grill-Spector, R. Goebel, A Probabilistic Functional Atlas of Human Occipito-Temporal Visual Cortex. Cerebral Cortex 31, 603–619 (2021).

101. E. H. Silson, I. I. A. Groen, D. J. Kravitz, C. I. Baker, Evaluating the correspondence between face-, scene-, and object-selectivity and retinotopic organization within lateral occipitotemporal cortex. J. Vis. 16, 14–14 (2016).

102. R. M. Hutchison, J. C. Culham, S. Everling, J. R. Flanagan, J. P. Gallivan, Distinct and distributed functional connectivity patterns across cortex reflect the domain-specific constraints of object, face, scene, body, and tool category-selective modules in the ventral visual pathway. Neuroimage 96, 216–236 (2014).

103. W. D. Stevens, M. H. Tessler, C. S. Peng, A. Martin, Functional connectivity constrains the category-related organization of human ventral occipitotemporal cortex. Hum. Brain Mapp. 36, 2187–2206 (2015).

104. E. H. Silson, A. D. Steel, C. I. Baker, Scene-Selectivity and Retinotopy in Medial Parietal Cortex. Front. Hum. Neurosci. 10, 412 (2016).

105. C. Baldassano, A. Esteva, L. Fei-Fei, D. M. Beck, Two Distinct Scene-Processing Networks Connecting Vision and Memory. eNeuro 3 (2016).

106. D. D. Dilks, J. B. Julian, A. M. Paunov, N. Kanwisher, The occipital place area is causally and selectively involved in scene perception. Journal of Neuroscience 33, 1331–1336 (2013).

107. U. Hasson, M. Harel, I. Levy, R. Malach, Large-scale mirror-symmetry organization of human occipito-temporal object areas. Neuron 37, 1027–1041 (2003).

108. N. Kanwisher, J. McDermott, M. M. Chun, The fusiform face area: A module in human extrastriate cortex specialized for face perception. Journal of Neuroscience 17, 4302–4311 (1997).

109. E. Halgren, et al., Location of human face-selective cortex with respect to retinotopic areas. Hum. Brain Mapp. 7, 29 (1999).

